# An ex vivo model of *Toxoplasma* recrudescence

**DOI:** 10.1101/2020.05.18.101931

**Authors:** Amber L. Goerner, Edward A. Vizcarra, David D. Hong, Kristina V. Bergersen, Carmelo A. Alvarez, Michael A. Talavera, Emma H. Wilson, Michael W. White

**Affiliations:** Department of Internal Medicine, Division of Infectious Disease and International Medicine, Morsani College of Medicine, University of South Florida, Tampa, Florida, USA; Division of Biomedical Sciences, School of Medicine, University of California, Riverside, Riverside, California, USA

**Keywords:** Apicomplexa, *Toxoplasma gondii*, development, tissue cyst, latency, chronic toxoplasmosis

## Abstract

*Toxoplasma* has been a useful parasite model for decades because it is relatively easy to genetically modify and culture, however, attempts to generate and study the recrudescence of tissue cysts have come up short with lab-adapted strains generating low numbers of tissue cysts in vivo. Here we have established a new model of *Toxoplasma* recrudescence using bradyzoites from an unadapted Type II ME49 strain (ME49EW) isolated from murine brain tissue. Ex vivo bradyzoite infection of fibroblasts and astrocytes produced sequential tachyzoite growth stages; a fast-growing stage was followed by formation of a slower-growing stage. In astrocytes, but not in fibroblasts, bradyzoites also initiated a second recrudescent pathway involving bradyzoite to bradyzoite replication. Intraperitoneal infections of mice with either bradyzoites or the fast-growing tachyzoite stage efficiently disseminated to brain tissue leading to high numbers of tissue cysts, while infections with the slow-growing tachyzoite stage were largely retained in the peritoneum. Poor infection and cyst formation of slow-growing tachyzoites was reversible by serial tissue cyst passage, while the poor tissue cyst formation of lab-adapted tachyzoites was not reversible by these approaches. To distinguish strain developmental competency, we identified *Toxoplasma* genes highly expressed in ME49EW in vivo tissue cysts and developed a qPCR approach that differentiates immature from mature bradyzoites. In summary, the results presented describe a new ex vivo bradyzoite recrudescence model that fully captures the growth and developmental processes during toxoplasmosis reactivation in vivo opening the door to the further study of these important features of the *Toxoplasma* intermediate life cycle.

## INTRODUCTION

Reactivation of the *Toxoplasma* tissue cyst is a significant health threat to people chronically infected with this parasite and is life-threatening to infected individuals that are or become immunocompromised. Individuals acquire their initial *Toxoplasma* infection by ingesting oocysts from environmental sources or tissue cysts from contaminated food products. Oocysts and tissue cysts are exposed to proteases in the GI track that release the sporozoite and bradyzoite stages, respectively, to invade epithelial cells lining the gut. The time from localized infection to systemic escape is relatively short in mice. By 2h post-oral oocyst infection, *Toxoplasma* parasites have infected the intestinal epithelium and by 8h the infection could be detected in the mesenteric lymph nodes (1). Based on sensitive bioassay methods, the tissue cyst form can be detected in 6-7 days in the brain of mice orally infected with the oocyst stage (2). The short timeframe from initial infection to the establishment of chronic disease makes it difficult, if not impossible, to prevent *Toxoplasma* infection, which accounts for the widespread distribution of this parasite. It is estimated that one third of human populations are infected with this pathogen (3). Healthy individuals are at low risk for clinical complications from toxoplasmosis, although 10-20% of immune competent adults and children are symptomatic for this disease (4). *Toxoplasma* is one of the most frequent causes of inflammation of the uvea layer of the eye (3) and in some cases these infections result in the loss of vision. Relapsing disease is common in the first two years following a primary lesion diagnosis, especially in elderly patients (5). Prior to the introduction of HAART-therapy, *Toxoplasma* caused frequent life-threatening encephalitis in AIDS patients, with reactivating disease thought to be the most common cause (3). Today, *Toxoplasma* infections of AIDS patients remain a significant clinical threat to large populations that have poor access to HIV therapies and this will remain so until there is a HIV cure and/or a therapy is found to eliminate the chronic tissue cyst stage.

Recrudescence of the *Toxoplasma* bradyzoite tissue cyst is central to reactivation of toxoplasmosis, which is a developmental process infrequently studied and poorly understood. Studies of severely immunosuppressed mice indicate that *Toxoplasma* reactivation preferentially occurs in the brain frontal and parietal cortex grey matter (6), which is composed of neuronal cell bodies, dendrites and unmyelinated axons, and astrocytes and oligodendrocytes. There were few reactivating lesions observed in the cerebellum indicating this region has few tissue cysts (6), which was confirmed by tissue cyst mapping in the infected murine brain (7). Attempts to define the parasite developmental process associated with recrudescence in immunosuppressed mouse models and in AIDS patients that have succumbed to toxoplasmosis have uncovered heterologous parasite development in foci of reactivation (8, 9). Many vacuoles of different sizes filled with parasites expressing tachyzoite and/or bradyzoite markers are present simultaneously in these lesions. Consequently, the original cyst rupture could not be identified nor could the sequence of parasite developmental steps or host cell contributions be discerned (8, 9). These few studies point to the challenges in studying *Toxoplasma* reactivation, which currently is limited to animal models. What is critically needed is an in vitro model of bradyzoite reactivation that could accelerate efforts to understand these processes at the cell and molecular level.

In this paper, we describe a new ex vivo model of *Toxoplasma* recrudescence. The crucial foundation of this model is the use of a robust cyst-forming strain that has never been adapted to cell culture. We define two distinct pathways of bradyzoite recrudescence that are host cell-dependent. In addition, we introduce new markers that are able to differentiate bradyzoites based on developmental maturity.

## RESULTS

### Adaptation to cell culture leads to the loss of *Toxoplasma* developmental competency

One of the major challenges in studying the tissue cyst stage has been the preservation of developmental competency in experimental models especially cell culture systems. The Type II ME49 strain has been regularly used by many labs because it results in a consistent and high number of cysts in the brain. The ME49 strain used here we have designated as ME49EW as it has been exclusively maintained in vivo by alternating passage through resistant and sensitive mouse strains in order to sustain high tissue cyst production for over 20 years (see Fig. S1 for maintenance practice). From an i.p. infection of 10 cysts, the native ME49EW strain consistently produced thousands of tissue cysts (cyst to cyst) in multiple mouse strains (CBA/j mouse EW infection, Fig. 1A). To demonstrate the problems associated with cell culture adaptation, we infected human foreskin fibroblasts (HFF) with ME49EW bradyzoites and after 6 attempts (∼30 million total bradyzoites plated) an HFF-adapted parasite line emerged that grew continuously in HFF cells under standard conditions (21% oxygen) and was cloned (ME49EW1 strain). An infection of 10,000 ME49EW1 tachyzoites was well tolerated by CBA/j mice, however, the competency to produce tissue cysts in murine brain tissue was dramatically reduced (EW1 tachy to cyst, Fig. 1A). Although we did not conduct a full kinetic analysis of tissue cysts size as was done previously (10), ME49EW cyst size at 42 days post-infection (PI) fell in a similar range with an average cyst diameter of 40μm as opposed to 30μm for ME49EW1 (Fig. 1B). A third of ME49EW cysts were >50 μm as compared to 13% of ME49EW1 cysts and we detected no ME49EW cysts <15 μm, which were common in ME49EW1 infections. The reduction of ME49EW1 cyst production could not be reversed by infections of mice with a standard 10 cyst (i.p.) dose of ME49EW1 tissue cysts recovered from the tachyzoite infected mice (EW1 cyst to cyst, Fig. 1A) and B1 gene-analysis verified that low ME49EW1 cyst counts in brain were not the result of a change in *Toxoplasma* distribution in mouse tissues (Goerner and White, data not shown). The loss of developmental competency following forced adaptation to HFF cell culture is not an isolated outcome (11). Infection of CBA/j mice with two common laboratory strains (10,000 tachyzoites, i.p.) used to study bradyzoite development in vitro (PruQ and ME49-Tir1) (12, 13), also produced far fewer tissue cysts in CBA/j mice as compared to native ME49EW parasites (Fig. 1A:PruQ=25 cysts; ME49-Tir1=150 cysts). In contrast to robust cyst wall formation in response to alkaline stress in cell culture, low numbers of in vivo tissue cyst numbers for the PruQ strain have been reported by many laboratories (recently cataloged in ref 14). This highlights the major limitation of the cyst wall as a functional developmental marker, which was recognized nearly 30 years ago (15) and even more recently (16). Similar to the ME49EW1 strain, secondary infections with PruQ tissue cysts did not reverse poor cyst production in CBA/j mice (PruQ C-to-C, Fig. 1A). The PruQ strain was engineered to express GFP under the control of the bradyzoite-specific LDH2 promoter (13), and therefore, bradyzoites in these cysts can be distinguished from tachyzoites by their green fluorescence. Microscopic examination of live PruQ tissue cysts from 30 day CBA/j infected mice showed that PruQ cysts had major gaps in GFP fluorescence (LDH2-GFP image, Fig. 1C), and these gaps contained parasites as shown by live DNA staining with Hoechst dye (Fig. 1C) indicating there is a mixture of tachyzoites and bradyzoites in these cysts. Unfortunately, there were too few PruQ cysts to extend the analysis to free bradyzoite populations. By comparison, ME49EW tissue cysts were typically larger at 30 day PI (Fig. 1C) and IFA analysis of purified ME49EW bradyzoites (Fig. S2B) determined they were all mature bradyzoites (no SAG1 expression). Cell cycle analysis of ME49EW bradyzoites showed they were >98% 1N genomic content (G1/G0)(Fig. S2A) and IFA analysis determined there were no mitotic forms present in over 1,000 bradyzoites examined (Fig. S2B, no u-shaped or double nuclei per parasite or internal daughter buds). The nuclei in ME49EW dormant bradyzoites had an unusual extreme posterior location (see inset image, Fig. S2B) likely due to the large number of amylopectin granules (see bradyzoite EM studies ref. 17) that displace the nucleus. In replicating tachyzoites, the nucleus is central (18) due to the physical constraints linking mitosis with daughter budding (19), and thus, it is not clear the far posterior location of dormant bradyzoite nuclei would be compatible with endodyogenic replication. Combined these results are consistent with a G1/G0 cell cycle status for ME49EW bradyzoites that is similar to Type III, VEG strain bradyzoites produced by sporozoite infections of CD-1 mice (20). In the absence of a suitable laboratory strain to investigate bradyzoite recrudescence, we optimized ME49EW tissue cyst production in CBA/j mice to reliably produce 5-10,000 tissue cysts per mouse cortex (cerebellum contains <5% of total brain cysts) with yields of 1-2 million purified bradyzoites (30-60% recovery) per mouse (see Materials and Methods and Fig. S1 for details)

**Figure S1.**
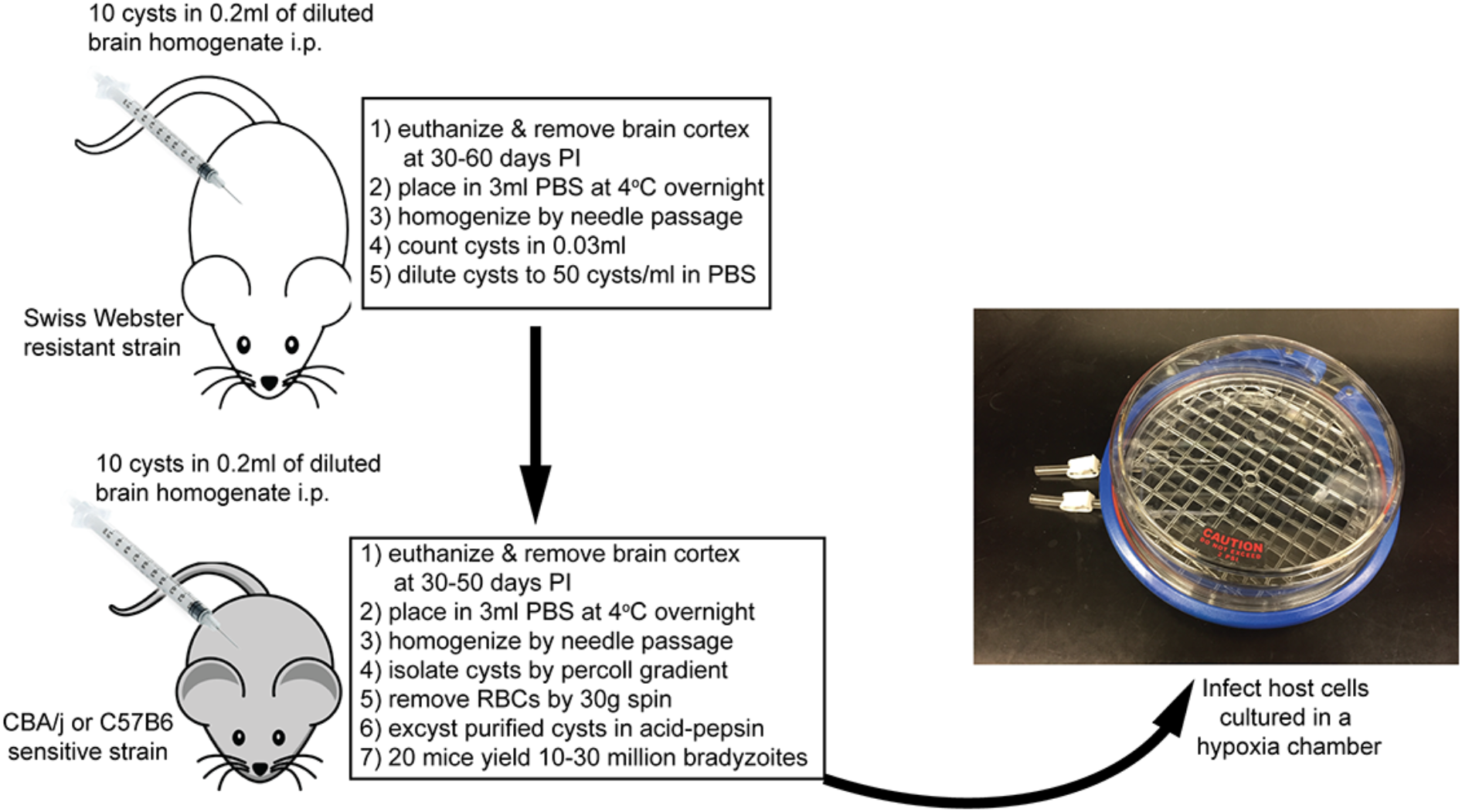
ME49EW tissue cyst maintenance and production. The ME49EW strain is maintained in the relatively resistant Swiss Webster (SWR/j) mouse strain by a 10 cyst inoculation (i.p.) every 30-60 days. Tissue cysts from Swiss Webster brain homogenates are used to amplify tissue cysts numbers in the sensitive CBA/j mouse strain (30-50 day infections). Sequential passage in CBA/j mice alone results in a progressive decline in tissue cyst number after multiple passages. The major steps for maintenance and experimental procedures are indicated with full details in Materials and Methods. Native bradyzoites in mouse brain reside in a hypoxic tissue environment, and as a consequence ME49EW strain bradyzoite-infected host cell monolayers are healthier if cultured at 37°C under 5% oxygen in HEPES buffered DMEM media containing 5% FBS (media recipe below). To achieve hypoxia conditions we use a hypoxia chamber (see picture) charged with mixed gas 90% nitrogen/5% oxygen/5% CO2 for 3 min and placed on the shelf of a CO2 incubator at 37°C. Using a multi-gas incubator commonly used to propagate *Plasmodium falciparum* merozoites in RBCs would also work. [Hypoxia chambers, https://www.stemcell.com/hypoxia-incubator-chamber.html] Growth media recipe for HFF and neonatal cells: Volume: 500ml --Use powder DMEM media with L-glutamine and sodium pyruvate (6.74g), --add 12.5ml 1M HEPES buffer, 25ml FBS, and 5 ml of antibiotic mix (Pen-strep and amphotericin B). --For astrocyte media, add 1% GIutaMAX supplement --Make the full media, then bring up to 37°C and adjust pH to 7.4 using 4M KOH. --Sterile filter.

**Figure 1.**
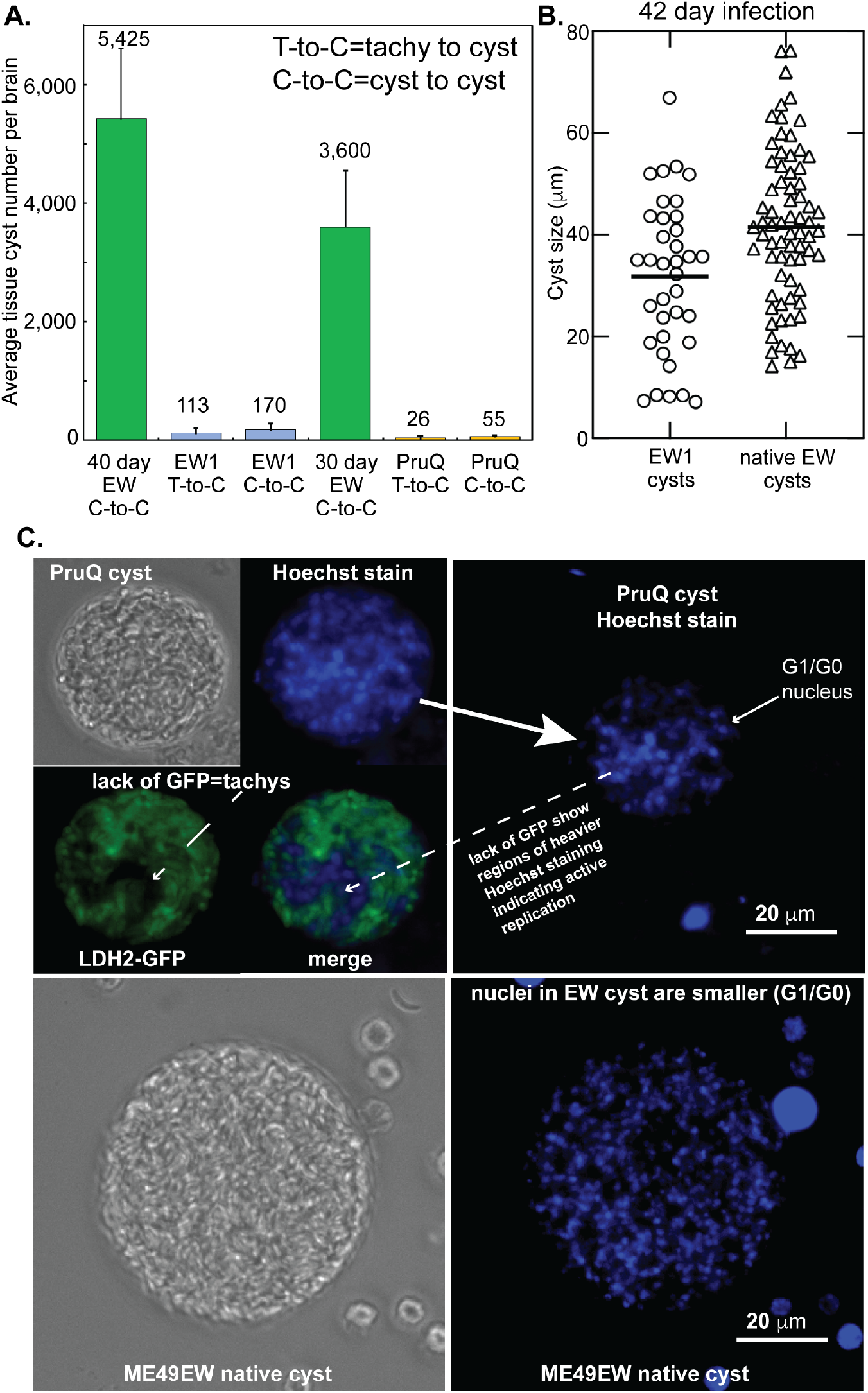
Adaptation to growth in HFF cells leads to a permanent loss of in vivo developmental competency. **[A.]** Type II strains, ME49EW, maintained in mice (see Fig. S1), an HFF-adapted strain derived from this strain, ME49EW1, and PruQ strain (PruQ=Prugnauid*Δku80Δhxgprt* ::LDH2-GFP) were used to infect six groups of 5 CBA/j mice. **Infections**: two groups of 5 mice were infected with ME49EW tissue cysts (C-to-C) and tissue cyst counts determined in brain tissue at 30 or 40 days PI; two groups of 5 mice were infected with ME49EW1 or PruQ tachyzoites (10k tachyzoites/mouse i.p.) and cyst counts determined at 30 day (PruQ, T-to-C) or 40 day (EW1, T-to-C); finally, EW1 or PruQ cysts (10 cysts, i.p.) from the T-to-C infected mice were used to infect two groups of 5 mice and cyst numbers determined at 30 day (PruQ, C-to-C) or 40 day (EW1, C-to-C). Brain tissue homogenates (3 ml) were prepared and total tissue cysts in 0.030 ml aliquots (1/100th of homogenate) were enumerated in at least triplicate by direct microscopic examination. Numbers above each bar=average cyst number. Note: A consistent 1.5 fold increase in tissue cyst number is observed between 30-40 days PI in ME49EW infections. **[B.]** Tissue cyst sizes from mouse brain tissue (42 day PI) were determined for native ME49EW and the lab-adapted strain, ME49EW1. **[C.]** Representative DIC and fluorescence images of live tissue cysts stained with Hoechst dye (top image row=PruQ and bottom row=ME49EW. All images are 400x magnification and were captured using the same mode fluorescent settings. Green=live bradyzoites (tachyzoites do not fluoresce), Blue=live Hoechst staining of genomic DNA. Note that PruQ cysts are typically smaller than ME49EW cysts at 30 day PI and contain large gaps in GFP fluorescence. The right hand Hoechst stained images for PruQ and ME49EW were adjusted identically to reduce fluorescent signal spreading in order to better capture nuclei size and relative DNA content. Note the small nuclei consistent with G1/G0 parasites are uniform and numerous in the ME49EW cyst. The nuclei in the PruQ cyst are nonuniform with large nuclei staining more intensely and co-localized with the gaps in the LDH2-GFP fluorescence suggesting these are replicating tachyzoites. Smaller nuclei are associated with the regions of GFP fluorescence in the PruQ cyst, which would be consistent with dormant bradyzoites.

**Figure S2.**
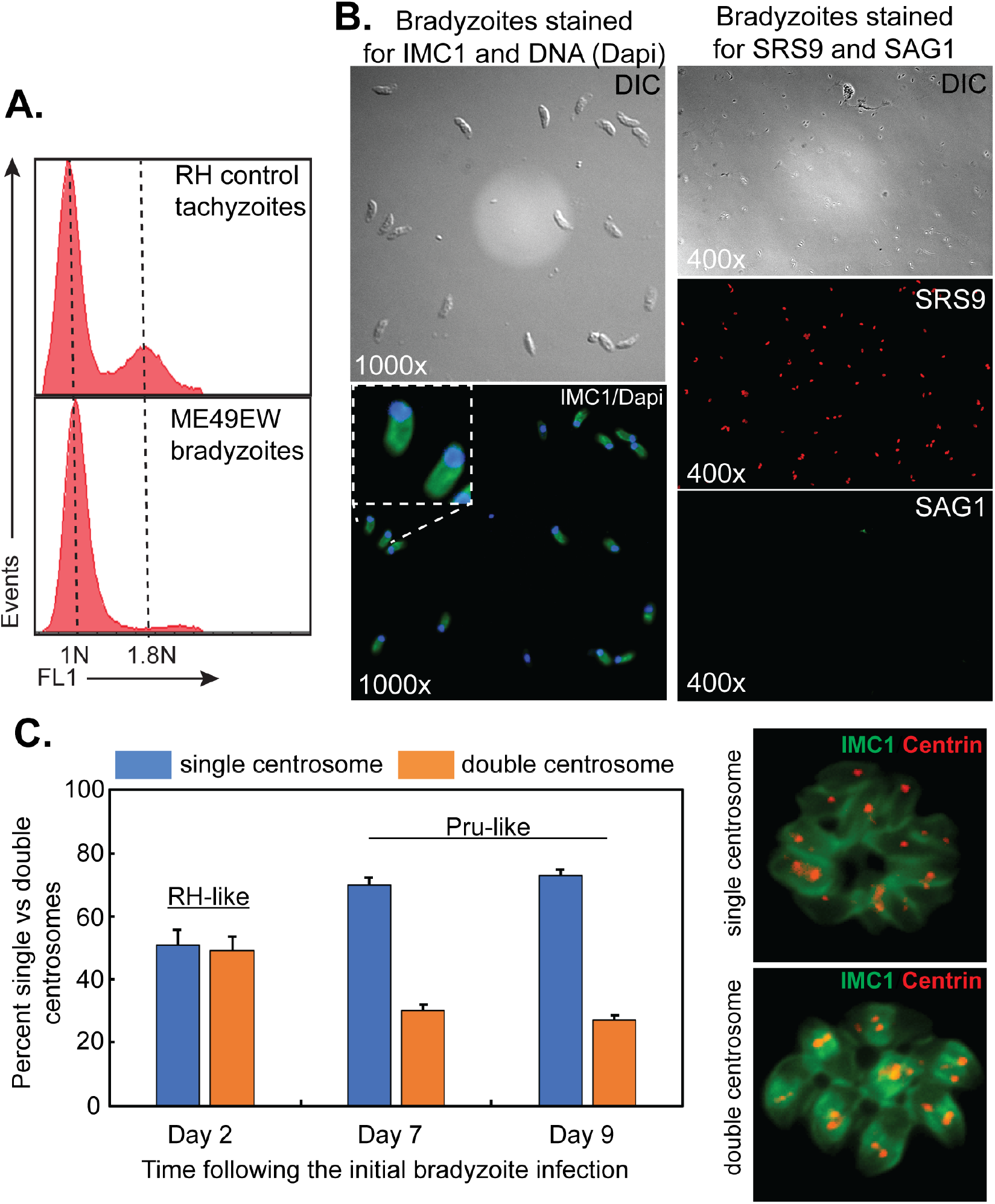
Cell cycle analysis of ME49EW parasite populations. **[A.]** Genomic DNA analysis of ME49EW bradyzoites in comparison to RH tachyzoites. Parasites were purified, ethanol fixed and stained with SYTOX dye and analyzed by flow cytometry. The cytometer was set to linear mode fluorescence and calibrated to the asynchronous RH control parasites; dashed lines reference 1N and 1.8N fluorescence peaks in the asynchronous control. DNA fluorescence was measured in FL-1 (x-axis) and 70,000 events (y-axis) were collected for each histogram. **[B.]**. Representative images of ME49EW bradyzoites co-stained for IMC1 (budding) and Dapi (nuclei) or separately co-stained for SRS9 (bradyzoite stain) and SAG1 (tachyzoite stain). Note all bradyzoites were SRS9+/SAG1-. Co-staining with an antibody against IMC1 and Dapi, showed ME49EW bradyzoites lacked internal daughters or u-shaped or duplicated nuclei demonstrating they were not replicating. In the vast majority of the bradyzoites the nucleus was in a far posterior location (see inset image) with very few bradyzoites having the central location of the nucleus in replicating tachyzoites (18). The IMC1-Dapi merge image shown is representative of ∼1,000 bradyzoites examined. **[C.]** At Day 2 (first monolayer), Day 7 (3rd monolayer), and Day 9 (4th monolayer) from the initial bradyzoite infection, parasites were fixed and co-stained for anti-centrin (centrosome), Dapi (DNA), and anti-IMC1 (internal daughters). Single versus double centrosomes were quantified in 50×3 vacuoles selected at random. The merged images (IMC1 and Centrin co-staining) to the right are representative of parasites possessing single versus double centrosomes.

### ME49EW bradyzoite infections initiate dynamic growth changes

In order to establish an ex vivo model of bradyzoite recrudescence, we investigated the developmental pathway that unfolds in ME49EW bradyzoite infections of two principle host cells, HFF cells and primary mouse neonatal astrocytes. The HFF host cell has been used to study *Toxoplasma* development for over 100 years and has allowed the standardization of techniques such as plaquing to be used to monitor growth. Astrocytes, which are relatively easy to culture, are a major host cell that bradyzoites encounter during recrudescence in brain and eye tissue (5, 21, 22). Initial ME49EW bradyzoite infections of HFF cells using standard tachyzoite culture conditions showed poor growth, which we reasoned might be due to the oxygen levels in cell culture as bradyzoites in mouse brain reside in a low oxygen environment (23). By lowering oxygen levels from 21% to 5% (see Fig. S1 for further details), we were able to establish more robust ME49EW parasite cultures in HFF cells. Lower oxygen levels did not harm HFF or astrocyte host cells nor did lower oxygen alter the growth of HFF-adapted parasite strains, and therefore, all ME49EW cell culture experiments were performed under 5% oxygen. To avoid host cells becoming a limiting factor and to allow for continuous tracking of the growth rate, parasites were passed after Day 3 and Day 5 (and sometimes after Day 7). This strategy is similar to the approach used to successfully characterize *Toxoplasma* sporozoite-initiated development (24) and is diagrammed in Figure 2A.

**Figure 2.**
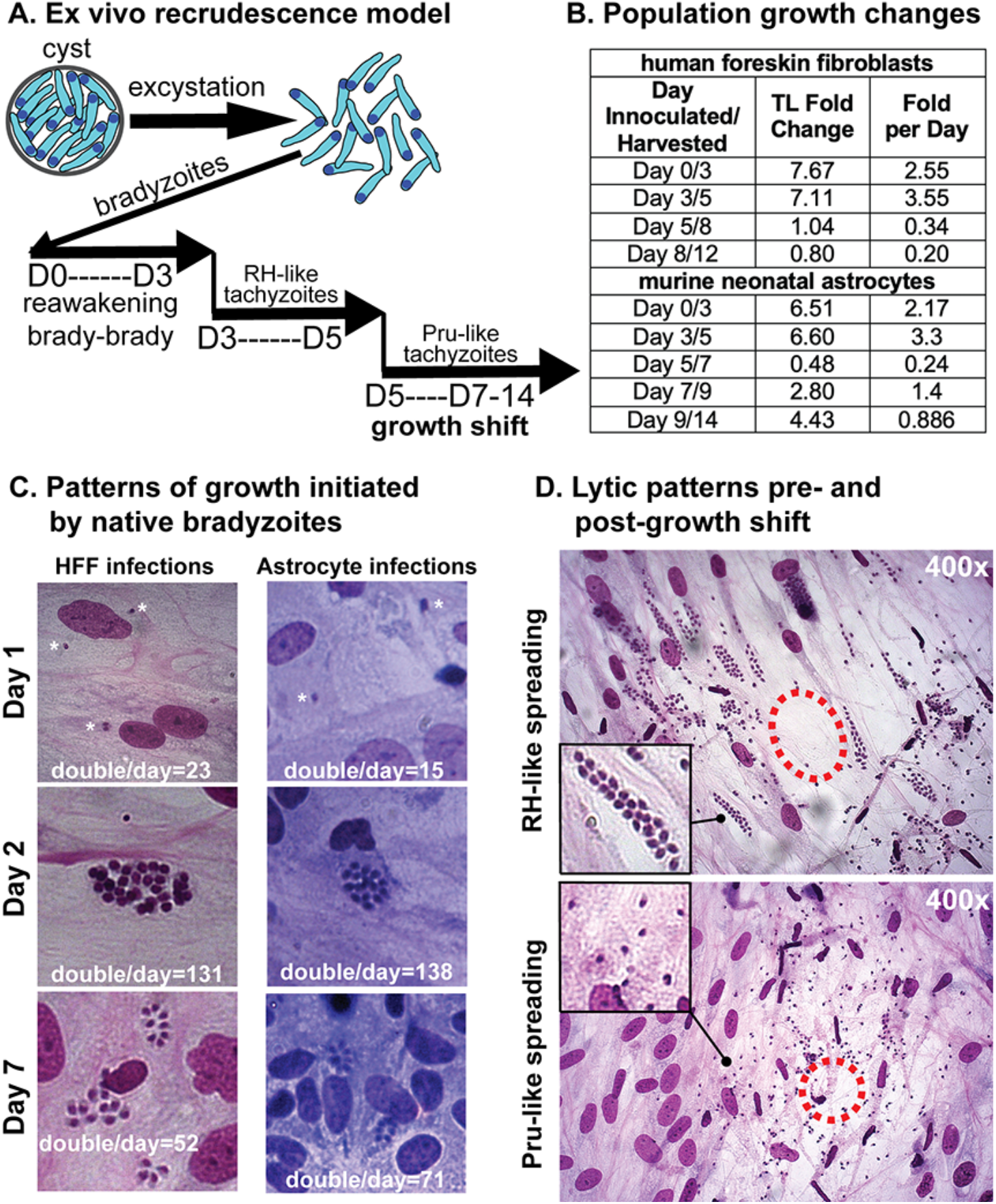
Native ME49EW bradyzoites initiate dynamic development. ME49EW bradyzoites from CBA/j murine brain tissue cysts were purified and used to inoculate monolayers of HFF cells or primary murine astrocytes (see Materials and Methods). **[A.]** Parasite populations are followed for 1-2 weeks and are labeled by the cumulative time from the original bradyzoite infection (D0). Day 3-populations are the end point of the cultures infected with bradyzoites; Day 5-populations are the end point of the cultures infected with Day 3 parasites; Day 7-populations are the end point of cultures of infected with Day 5 parasites etc. **[B.]** Representative total (TL) population growth during each growth period in HFF or astrocyte monolayers. The growth periods ranged from 2 to 5 days as indicated and Day 0 corresponds to the starting bradyzoite infection. **[C. and D.]** At various times, H&E stains (nuclei stain dark blue to violet) were prepared and vacuole size and lytic patterns were evaluated. Representative images of the major vacuole sizes. Total doublings (e.g. vacuole of 32=5 doublings) in 50 randomly selected vacuoles were determined in triplicate and the average parasite doublings per day are indicated at the bottom of each image. Note: Day 1-populations showed minimal growth (astrocytes=70% singles; HFF=60% singles), therefore, we subtracted Day 1 doublings from Day 2-population values in order to estimate the average doublings in the 24h period from the end of Day 1 to the end of Day 2. Asterisks in the top two images indicate the single bradyzoite vacuoles. Representative lytic patterns of Day 3- and Day 11-populations in HFF cells (first and fourth growth periods). Dashed circles indicate the approximate center of each plaque. Inset images=higher magnification of equal areas of the plaque, note the nuclei size differences and relative MOI differences in each plaque.

Purified bradyzoites from ME49EW tissue cysts readily invaded HFF cells and astrocytes and followed a similar course of growth in both host cells during the first week. At the end of the first 24h in HFF or astrocytes (Day 1-populations) the majority of vacuoles contained single parasites resulting in relatively few parasite doublings during this growth period (Fig. 2C, Day 1). Parasite replication quickly accelerated in the second 24h period (Day 2-populations) with many vacuoles containing 16 or 32 parasites by the end of Day 2 (doublings increased >5 fold, Fig. 2C). We estimated that ME49EW parasites post-Day 1 were dividing at rates equivalent to the fastest growing laboratory strains (e.g. 6-8h doubling of RH tachyzoites). Parasites growing at these faster rates possessed a larger nucleus, invaded nearby host cells at high MOI and the plaques showed a relatively clear lysis zone with less cellular debris (Fig. 2D, RH-like spreading). We have designated this type of growth as “RH-like”. ME49EW population growth was similar in the first two growth periods (0-3 days and 3-5 days) in both host cell types (Fig. 2B). One week from the initial bradyzoite infection, parasites in either host cell type had spontaneously slowed as evident by smaller vacuole sizes and lower doubling rates in the third 48h growth period (Day 7-images, Fig. 2C). Spreading behavior was also changed particularly in HFF cells with plaque development similar to many slow-growing laboratory strains; far less multiple invasions of nearby cells, parasites with smaller nuclei (indicative of G1), and cellular debris in a less defined lysis zone (Fig. 2D, Pru-like spreading). We have designated this type of growth “Pru-like”. Longer cell cycle times in *Toxoplasma* laboratory strains are the result of extended G1 periods (18), which can be estimated from centrosome patterns. ME49EW parasites possessing single (G1 phase) versus double (S and M/C phases) centrosomes were equal in RH-like populations and nearly 2:1 in Pru-like populations (Fig. S2C). Slower growth became the dominant phenotype of all ME49EW populations post-Day 7 (Fig. 2B) and we never observed Pru-like parasites reverting to RH-like growth in HFF or astrocyte cell cultures. Beyond one week, ME49EW parasites in HFF cells suffered a continued productivity decline (Fig. 2B) and became difficult to sustain. Second week cultures of ME49EW parasites in primary astrocytes continued to grow at a rate half that of Day 2-5 parasites, however, production did not further decline in astrocytes through Day 14 (Fig. 2B).

To better understand the nature of ME49EW parasite growth in HFF and astrocytes, we assessed the extracellular viability of parasites produced by bradyzoite infections in each host cell type. Rapidly growing RH-like parasites from Day 2-populations in HFF cells or astrocytes were purified and resuspended in culture media. The extracellular parasites were sampled 0, 6, 12 and 24h and inoculated into either HFF or astrocyte monolayers. Plaque formation was quantified 4 days later by IFA assay (Fig. S3A, only 0, 6, and 12h timepoints shown). Native ME49EW Day 2-parasites from either host cell type were less stable in media with ∼88% of the parasites dying by 6h (2-3h half-life). By contrast, the HFF-adapted RH strain showed the opposite phenotype; extracellular incubation for >12h in media was required before ∼90% of RH parasites died. We next examined extracellular stability of the Type I strain, GT1 (25), as well as the ME49EW1 strain, which are low passage HFF-adapted clones. ME49EW1 and GT1 tachyzoites showed intermediate extracellular half-life as compared to native ME49EW Day 2-parasites or RH parasites (Fig. S3A). Taken together, these results are a caution that native *Toxoplasma* strains may have lower stability in culture media and must quickly enter host cells to survive, whereas adapting strains to growth in HFF progressively selects for increased extracellular stability, which is related to stress resistance (26).

**Figure S3.**
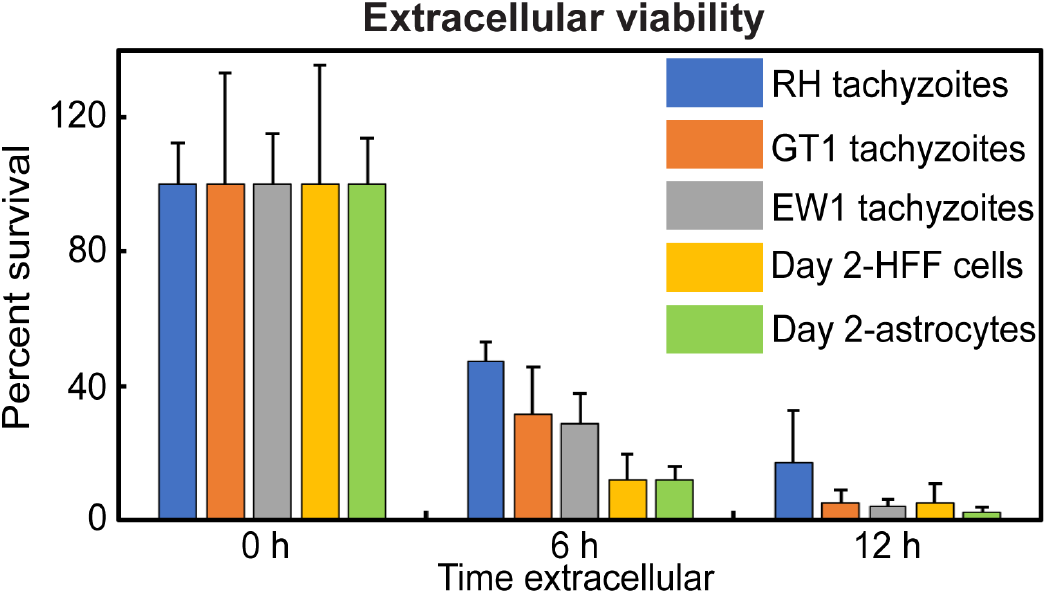
Extracellular half-life. Day 2-recrudescence populations from ME49EW bradyzoite infections of HFF or astrocyte monolayers and lab-adapted strains, Type II-ME49EW1 or Type I-RH and -GT1 strain tachyzoites were purified by standard methods and free parasites were resuspended at 1 million/ml of media and incubated at 37°C. At time zero, 6h, and 12h, 10,000 parasites were sampled and used to infect duplicate cover slips. After ∼4 days of further growth, cover slips were fixed and stained with goat, anti-*Toxoplasma* antibody. The number of plaques in 10 microscopic fields were determined in triplicate (6 total counts/timepoint). All results compared to time-zero.

### Host cells influence bradyzoite recrudescence

To understand the developmental changes associated with ME49EW bradyzoite recrudescence, we investigated the expression of two key surface antigens; SRS9 present on bradyzoites and the tachyzoite surface antigen SAG1 (Figs. 3,4 and S4,S5). As shown above (Fig. S2B), native bradyzoites invading host cells were uniformly SRS9+/SAG1- and this did not change for many hours after invasion (Fig. 3A). By the end of Day 1, most vacuoles in either host cell type had a single SRS9+/SAG1-parasite (70% in astrocytes and 60% in HFFs). Between Day 1 and Day 2 post-bradyzoite infection, SRS9+ expression began to decline as SAG1 expression increased with some vacuoles having equal expression of these antigens (astrocyte images, Fig. 3A; HFF images Fig. S4). The trend towards elevated SAG1 and low to absent SRS9 expression progressively became the dominant phenotype as seen in Day 3-population images (Fig. 3A, quantified in D), and especially Day 5-populations (Fig. 3A,B) that formed in either host cell type. For the first time in a cell culture model, we observed SRS9+ only bradyzoites replicating in the first astrocyte monolayer, while also not turning on SAG1 expression (Day 2, Fig. 3C, additional examples Fig. S5A). This bradyzoite replication occurred simultaneously to tachyzoite replication in neighboring cells (Figs. 3C and S5A). Bradyzoite replicating vacuoles were 15% of bradyzoite-infected astrocytes (Fig. 3D) and grew slower (Fig. 3E) progressing from an average vacuole size of 5.22 bradyzoites (SAG1+ tachyzoite vacuole=10) at Day 2 to a vacuole size of 12 bradyzoites by Day 3 (SAG1+ tachyzoite vacuole=21). Bradyzoite only replicating vacuoles (SRS9+/SAG1-) were observed in Day 5-populations (e.g. Fig. 4B), although at lower frequencies. The majority of Day 5-vacuoles were either SRS9+/SAG1+ or SRS9-/SAG1+ (Fig. 3A,B). SAG1+ only parasites continued to dominate populations in HFF cells and astrocytes even after the growth of parasites had spontaneously slowed. However, in astrocytes a small proportion of vacuoles (4%) containing SRS9+ only parasites were detectable after the Day 7-growth shift (see Fig. 4A for examples). While it was not possible to rule out that bradyzoite to bradyzoite replication gave rise to the Day 7-SRS9+/SAG1-vacuoles, there were many SRS9^Hi^/SAG1^low^ vacuoles (>30%) in Day 7-astrocyte cultures suggesting it was more likely parasites had invaded as SAG1+ parasites and then during replication turned back on SRS9 expression as SAG1 declined (tachy to brady development). In contrast, HFF cells did not support bradyzoite-to-bradyzoite replication. Only rare vacuoles staining exclusively for SRS9+ were present beyond Day 1, however this minor fraction of vacuoles contained single parasites (Fig. 3E). It is not known if these were nonreplicating bradyzoites being carried over or parasites that did not grow after passage and then re-expressed SRS9, while turning off SAG1.

**Figure 3.**
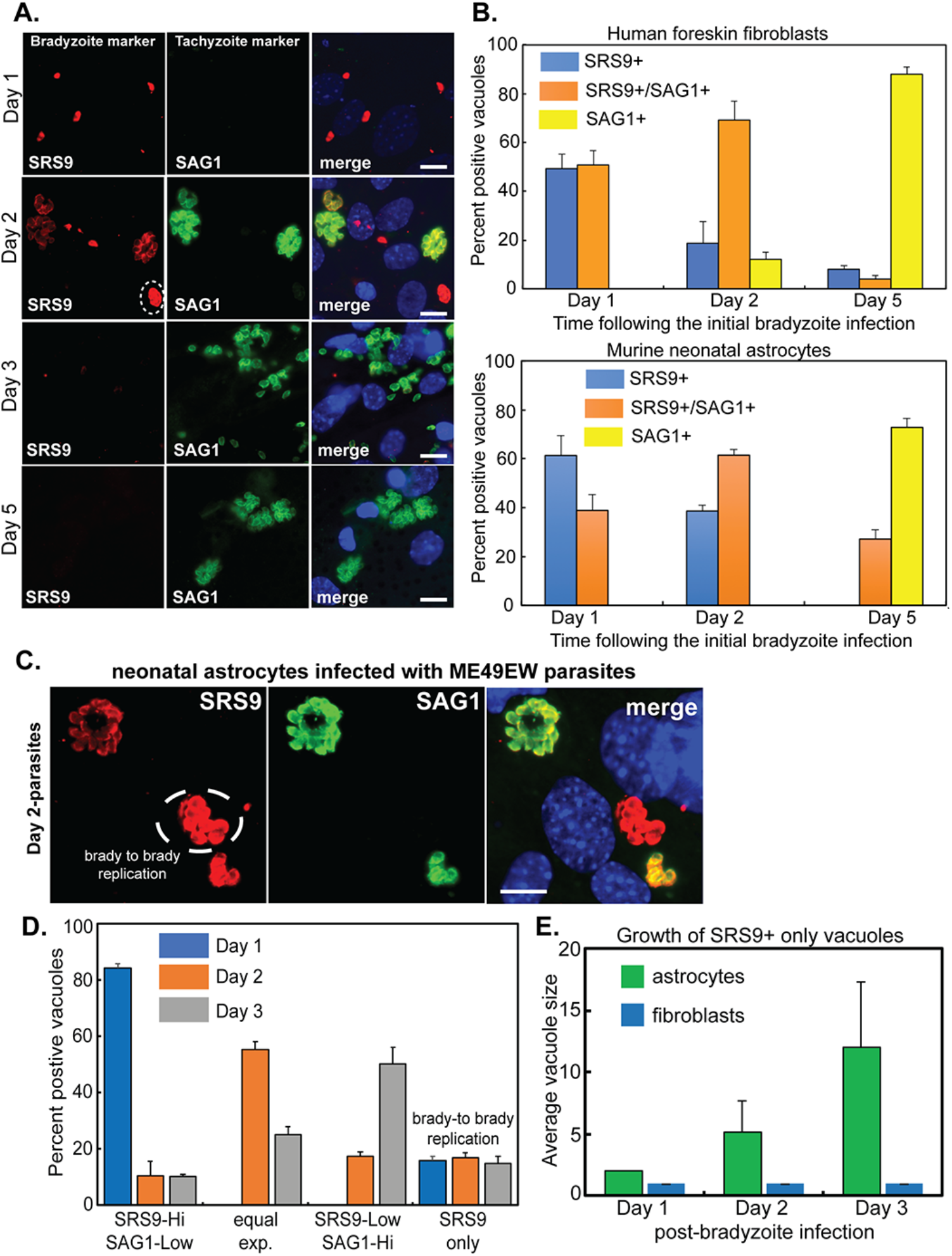
Changes in bradyzoite and tachyzoite surface antigen expression. SAG1 and SRS9 are developmentally regulated SRS antigens specific for tachyzoite and bradyzoite stages, respectfully. At various times following an initial bradyzoite infection of astrocytes or HFF cells IFA analysis was performed using co-stains for SAG1 (green), SRS9 (red) and Dapi (DNA, blue). (see Fig. S4 for infected HFF images). Note that parasites appear rounder in astrocytes than HFF cells due the thicker monolayer. **[A.]** Representative images of parasite staining patterns Day 1-3 post-infection of astrocytes with ME49EW bradyzoites. Day 5-images were of astrocyte monolayers infected with parasites purified from the Day 3-astrocyte culture and grown for 48h. Note the dashed circle in the Day 2-image is a SRS9-Hi/SAG1-low vacuole, while the other three replicating vacuoles in this microscope field are SRS9-low/SAG1-Hi. Scale bars=10 μm **[B.]** Parasite vacuoles in HFF cells or astrocytes expressing SAG1 and/or SRS9 were quantified at the indicated times. The SRS9-only vacuoles (blue bar) in HFF cells were single parasite vacuoles as there was no evidence of bradyzoite to bradyzoite replication observed in HFF cell IFAs (see E. below). **[C.]** Representative images of bradyzoite to bradyzoite replication (dashed circle) from a Day 2-astrocyte cell culture (vacuole of 8 parasites). For additional examples, see Figure S5A. Parasites were co-stained for SRS9 (red), SAG1 (green) and Dapi (merged image shows host cell nuclei). Scale bars=10 μm **[D. and E.]** Quantification of the developmental patterns of replicating vacuoles in bradyzoite-infected astrocyte monolayers and the average vacuole size of SRS9+/SAG1-vacuoles in HFF versus astrocytes.

**Figure S4.**
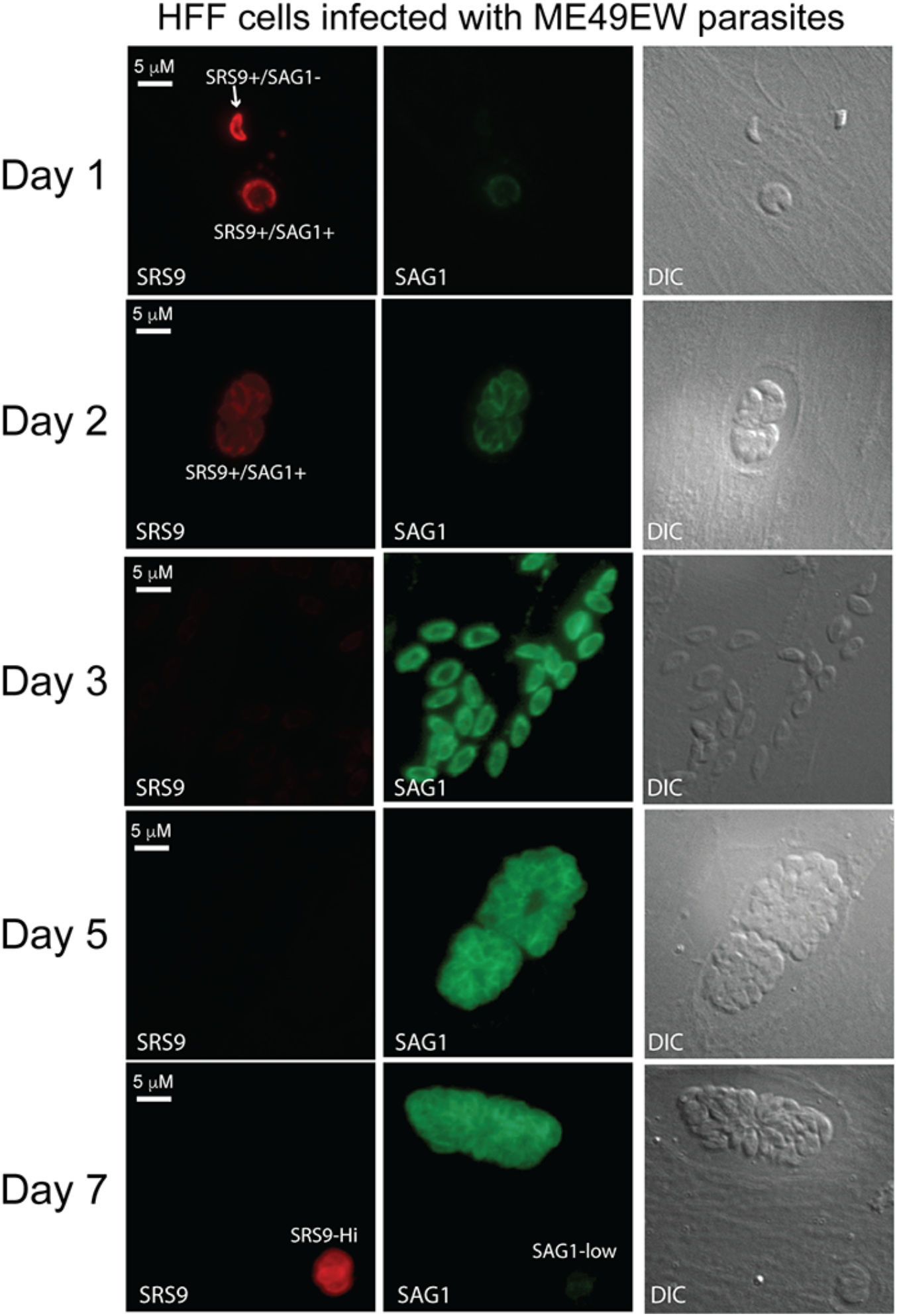
Changes in parasite developmental antigen expression in HFF cells. Representative images of parasite staining patterns Day 1-3 post-infection of HFF cells infected with native bradyzoites purified from mouse brain. Day 5 and Day 7 images were of HFF cells infected with parasites purified from Day 3- or Day 5-parasite populations and then grown for 48h to yield Day 5-images and Day 7-images, respectively. Parasites were fixed and co-stained for SAG1 (green), SRS9 (red). Paired DIC images are shown on the right.

**Figure 4.**
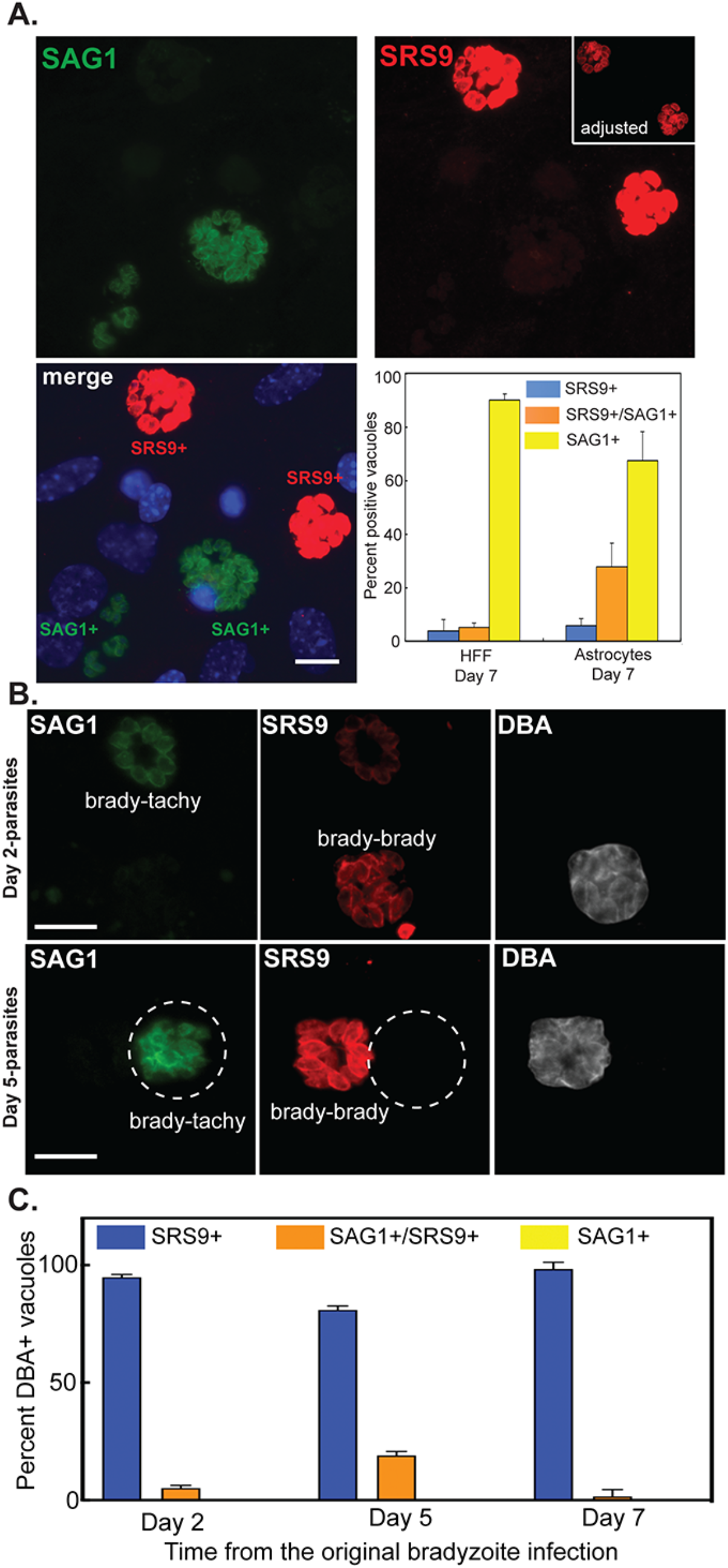
Post-growth shift development and assembly of cyst walls in astrocytes. **[A.]** SRS9 expression was assessed by IFA in post-growth shift infected astrocytes (Day 7-populations). Infected astrocytes were fixed and co-stained for SAG1 (green) and SRS9 (red) with Dapi staining included as a reference (DNA, blue). Representative images of SAG1+ or SRS9+ only parasite vacuoles in astrocyte-infected cells. Scale bars=10 μm Note two levels of SRS9 fluorescence are shown; the main images display the relative SRS9 (brighter signal) to SAG1 fluorescence signal, while the inset box displays a lower mode fluorescence for SRS9 that more clearly shows individual parasites in these vacuoles. Graph: quantification of SRS9 only, SRS9 & SAG1, or SAG1 only in Day 7-populations in HFF or astrocyte cells is also shown. Note that the Day 7-populations in HFF cells have higher levels of SAG1 only vacuoles. There are also rare SRS9 only vacuoles (blue bar) containing single parasites (∼1/5 of the 20% single parasite vacuoles in the HFF cultures). By contrast, there were no single parasite vacuoles observed in Day 7 astrocyte cultures. **[B and C.]**. Representative images and quantification of parasite vacuoles where cyst wall formation was detected. Parasites in astrocytes were co-stained with antibodies against SRS9 (red) and SAG1(green) and with biotin-labeled *Dolichos biflorus* agglutinin (DBA) to identify vacuoles possessing cyst walls. Size bars (10 μm) are indicated and the dashed circles indicate the vacuole containing SAG1+ parasites. Note: no vacuole containing SAG1+ only parasites showed evidence of cyst wall formation and only a minor fraction of vacuoles with parasites expressing both SRS9 and SAG1 were found to have cyst walls. Vacuoles with parasites expressing SAG1 only or both surface antigens representative of bradyzoite-to-tachyzoite recrudescence and vacuoles containing SRS9+ only parasites representing bradyzoite-to-bradyzoite replication are indicated.

**Figure S5.**
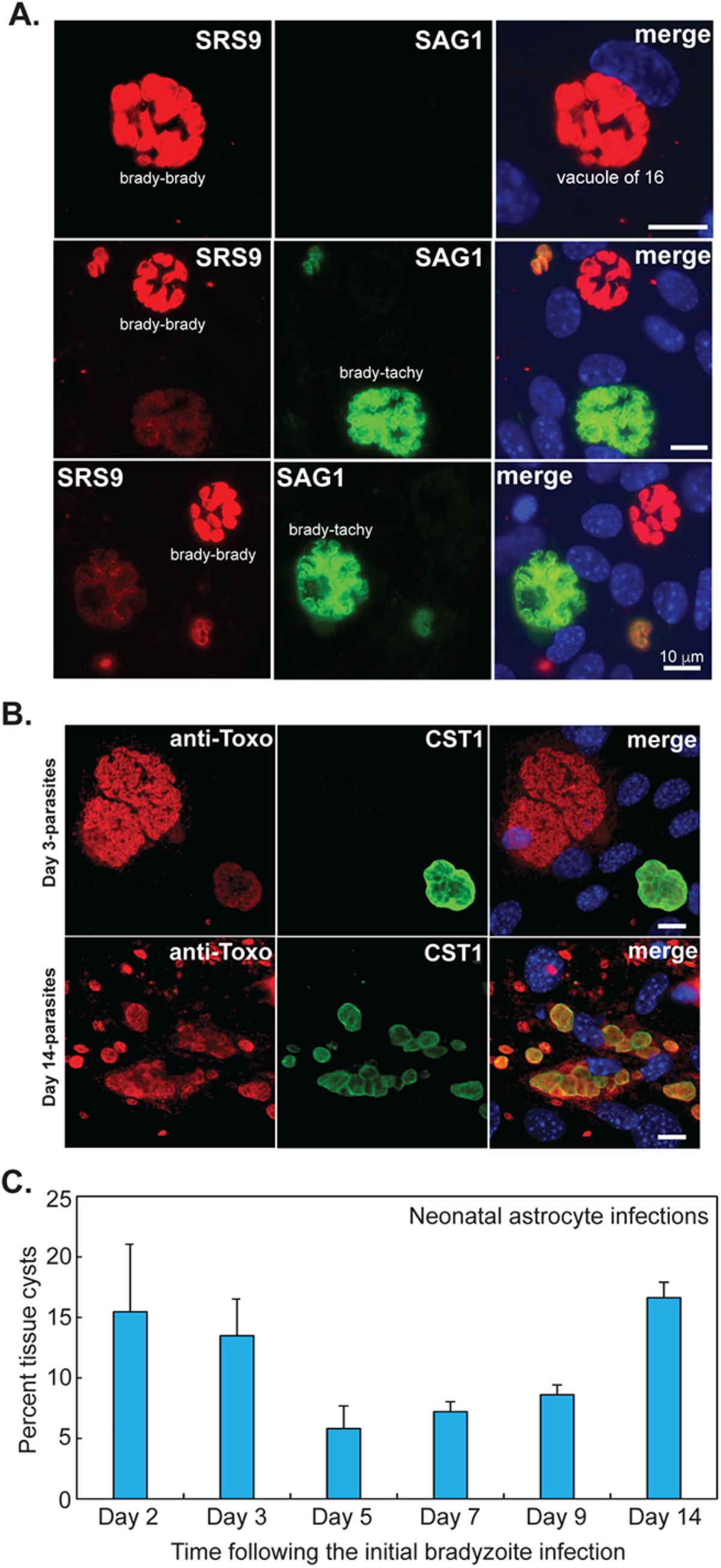
**[A.]** Additional examples of bradyzoite to bradyzoite replication (SRS9+/SAG1-) occurring along side bradyzoite to tachyzoite replication in astrocytes infected with ME49EW bradyzoites. merge image=co-staining for SRS9 and SAG1 and Dapi, which stains the large astrocyte nucleus. **[B and C.]** Representative images of tissue cysts in ME49EW infected astrocytes at Day 3 (first growth period) and Day 14 (fourth growth period) from the original ME49EW bradyzoite infection. CST1 staining (green), anti-*Toxoplasma* (red), and Dapi (blue). Scale bars= 10 μm. Note the Day 3-image shows a single cyst along side two large vacuoles of rapidly growing tachyzoites. The cluster of cysts in the Day 14 image highlights the focal nature of some cyst formation in these cultures. Graph: quantitation of the average percent of vacuoles containing a tissue cyst wall at various times from the original bradyzoite infection. Growth/monolayer periods: Day 0-3, Day 3-5, Day 5-7, Day 9-14.

We next investigated the spontaneous formation of tissue cysts during ME49EW recrudescence using lectin binding to the cyst wall (DBA+, Figure 4B and C) or antibodies against the cyst wall protein, CST1 (Fig. S5B). In astrocytes infected with bradyzoites (Days 1-3) spontaneous cyst wall formation occurred in 13-15% of vacuoles (CST1 expression, Fig. S5B and C) and those vacuoles were primarily undergoing bradyzoite to bradyzoite replication (Fig. 4B and C). This is strikingly different from lab-adapted tachyzoites induced to assemble cyst walls in alkaline-media, while also continuing to express high levels of SAG1 mRNA (see relative SAG1 mRNA levels in Fig. 5B) and SAG1 antigen (27). Cyst wall formation decreased to ∼5% in Day 5-populations (Fig. S5C) and then increased steadily following the shift to slower growth at Day 7 (Fig. S5B and C), although here again these new cyst walls were forming primarily in vacuoles containing replicating bradyzoites (Fig. 4B and C). By contrast in all infected HFF cells, spontaneous cyst wall formation occurred in <1% of the ME49EW vacuoles because in these experiments using the ME49EW strain there was little to no bradyzoite replication in this host cell type.

**Figure 5.**
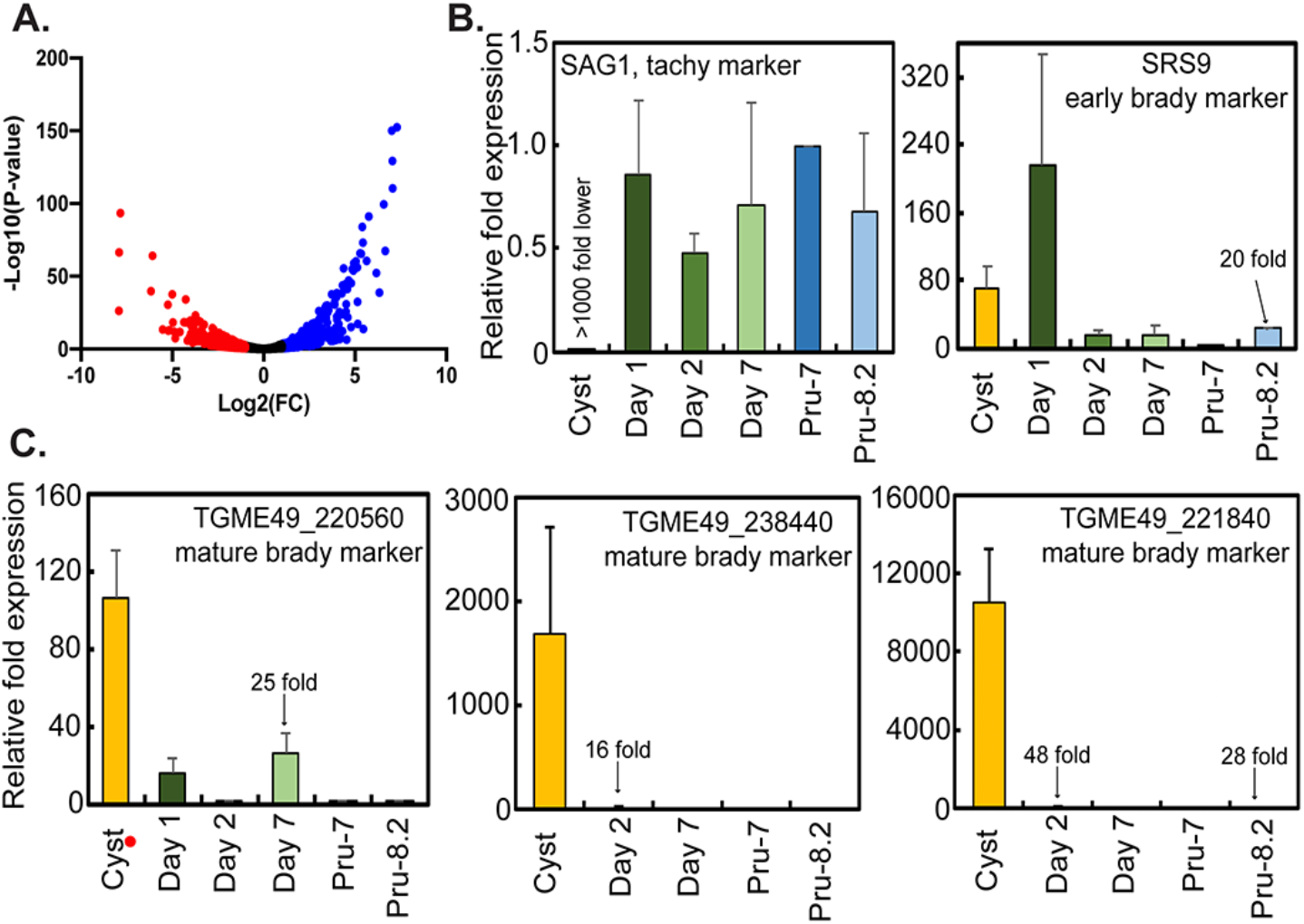
Bradyzoite gene expression in ME49EW in vivo tissue cysts. **[A.]** Analysis of whole-cell mRNA expression of Type II, ME49EW tissue cysts at 40 day post-infection in comparison to ME49EW1 tachyzoites. See Materials and Methods for full details. **[B. and C.]** qPCR gene expression analysis of SAG1 and SRS9 mRNAs in comparison to three in vivo-tissue cyst mRNAs. Results are presented as relative fold expression over the corresponding PruQ tachyzoite 2^-ΔCt^ values. RNA sources: ME49EW tissue cysts and Day 1-, 2-, and 7-recrudescent populations grown in HFF cells; PruQ strain tachyzoites (Pru-7) and tachyzoites induced to differentiate in pH8.2 media (Pru-8.2).

### Defining new markers of mature bradyzoites

It is clear that strains capable of robust in vivo cyst formation, such as ME49EW, are developmentally distinct from many HFF-adapted strains, although there are currently no molecular markers to distinguish these parasite strains. To identify new markers for the developmental competent ME49EW bradyzoite, we analyzed the transcriptome of ME49EW tissue cysts by RNA-sequencing (RNA-seq)(Fig. 5 and Dataset S1). Two independent samples of ME49EW tissue cysts were purified from mice that had been infected with 10 cysts i.p. and harvested at 40 day PI. Tissue cysts were lysed, RNA purified and RNA-libraries prepared and sequenced as described in Materials and Methods. Illumina sequencing of the two ME49EW cyst RNA-libraries yielded >50 million reads with 13.8% aligned to the *Toxoplasma* reference genome. We analyzed ME49EW tissue cyst transcriptome in comparison to the ME49EW1 tachyzoite transcriptome (Fig. 5A, volcano plot). Using a ≥2 fold cutoff, 2,263 mRNAs were differentially expressed in ME49EW cysts as compared to ME49EW1 tachyzoites; 933 mRNAs were upregulated, 1,330 mRNAs were downregulated (Fig. 5A and Dataset S1). The ME49EW tissue cyst transcriptome was also compared to the transcriptome of PruQ tachyzoites (see Dataset S1). Among the key gene expression differences between ME49EW tissue cysts and ME49EW1 and PruQ tachyzoites was the significant downregulation of many growth-related mRNAs (Dataset S1). Multiple DNA and RNA polymerases and DNA replication cofactors, 14 IMC proteins including the IMC1 marker of budding used here are markers for budding, mitotic factors such as centrin, as well as the classic S phase cell cycle marker dihydrofolate reductase-thymidylate synthase are all significantly downregulated in ME49EW tissue cysts. This transcriptional profile is consistent with the evidence shown earlier (Fig. S2B) that bradyzoites in these cysts are in a deep growth arrested state (G1/G0).

In order to identify mRNAs that could distinguish in vivo mature bradyzoites in practice, we first identified mRNAs highly expressed in ME49EW cysts as compared to ME49EW1 tachyzoites (>5 fold, Dataset S1). We then ranked the ME49EW bradyzoite-mRNAs based on abundance (>95 percentile) and finally, compared the list of ME49EW bradyzoite genes to normalized RNA-seq values from four datasets downloadable from ToxoDB; ME49 tachyzoites (36h), ME49 pH8.2 media-induced bradyzoites, as well as datasets derived from sporozoites and merozoites (see Dataset S1 gene list and dataset details). To develop a qPCR assay for mature bradyzoites, we selected three mRNAs (unknown-TGME49_220560, SRS22A-TGME49_238440, unknown-TGME49_221840) highly expressed in ME49EW cysts from mice, but expressed at much lower levels in alkaline-stressed ME49 parasites (Fig. 5C mature cyst markers). The features of this assay included primer sets for the tachyzoite marker SAG1, early bradyzoite marker SRS9, and actin depolymerizing factor as an internal reference gene. Each primer set passed standard quality control tests for use in qPCR. Total RNA was purified from ME49EW cysts and infected HFF monolayers of Day 1-, Day 2-, and Day 7-ME49EW recrudescent populations, and RNA was also purified from PruQ tachyzoites grown in normal pH 7.2 media or induced to differentiate in pH8.2 media (48h in vitro bradyzoites). Total RNA was converted to cDNA and used as template for SYBR-green qPCR analysis (see Dataset S1 for primers). A comparison of gene expression in these RNA sources defined two general states of bradyzoite differentiation (Fig. 5B,C). A pattern of increased early bradyzoite marker (SRS9), continued SAG1 expression and low to intermediate levels of mature bradyzoite markers indicated a state of bradyzoite immaturity, which was characteristic of the alkaline-media induced PruQ parasites and also ME49EW recrudescent tachyzoite RNA samples. By contrast, high levels of SRS9 and mature marker gene expression and the absence of SAG1 mRNA in ME49EW cyst RNA was consistent with bradyzoite maturity.

### Bradyzoites and recrudescing populations have distinct capacities to form tissue cysts in mice

To understand the biological function of the three growth phases of excysted parasites, we determined the cyst-forming capability of ME49EW dormant bradyzoites, RH-like and Pru-like populations in CBA/j mice. Bradyzoites were obtained by excystation of purified ME49EW tissue cysts, while RH-like parasites were purified from bradyzoite-infected HFF or astrocytes at Day 3 PI. Pru-like parasites were from Day 7 growth-shifted populations. For parasites grown in HFF cells, three groups of ten CBA/j mice were infected with a dose of 10,000 parasites i.p. and tissue cysts were quantified in groups of five mice at 14 and 30 days PI (Fig. 6A). For parasites grown in astrocytes, three groups of 5 CBA mice were infected and cysts counted at 30 days PI. Purified bradyzoites (Fig. 6A, closed circles) produced the highest number of brain cysts indicating intact cysts are not required to achieve robust cyst numbers in mouse brain. RH-like parasites (closed squares) also produced reasonable cyst numbers, although cysts counts from RH-like infections derived from HFF cells were not as high as the paired bradyzoite and RH-like populations derived from astrocytes. This difference could be experimental or biological and will require further investigation to resolve. Regardless of the host cell used, infections with Pru-like parasites resulted in significantly lower cyst numbers in CBA/j brain tissue. Unlike lab-adapted parasite strains, the reduction in cyst formation of ME49EW Pru-like parasites was not permanent as infections with 10 cysts from the first round of mouse infections gave equivalent cyst numbers irrespective of the source of the original parasite infection (Fig. 6B, 2nd round infections). Relative parasite burden in the brain of these mice correlated well with the cyst count data (Fig. 7). For example, in the day 14 PI group the mice infected with bradyzoites showed the highest B1 gene signals in brain tissue with RH-like and Pru-like parasites having stepwise lower burdens (Fig. 7 A,B). Other tissues had detectable B1 gene levels that were a fraction of the levels in brain indicating that changes in tissue distribution could not explain the cyst count differences. Because cytokine responses are related to parasite burden, we collected blood at various times in the mice scheduled for tissue cyst counts at Day 14 PI and quantified IFNγ levels (Fig. 8). Peak IFNγ levels seen at seven days post-infection with bradyzoites and RH-like parasites follows a timing previously reported (28), while IFNγ levels in Pru-like infections confirms successful infection and parasite replication but the reduction in IFNγ concentrations and delay in kinetics was consistent with the significantly lower parasite burden in mice infected with these parasites (Fig. 7).

**Figure 6.**
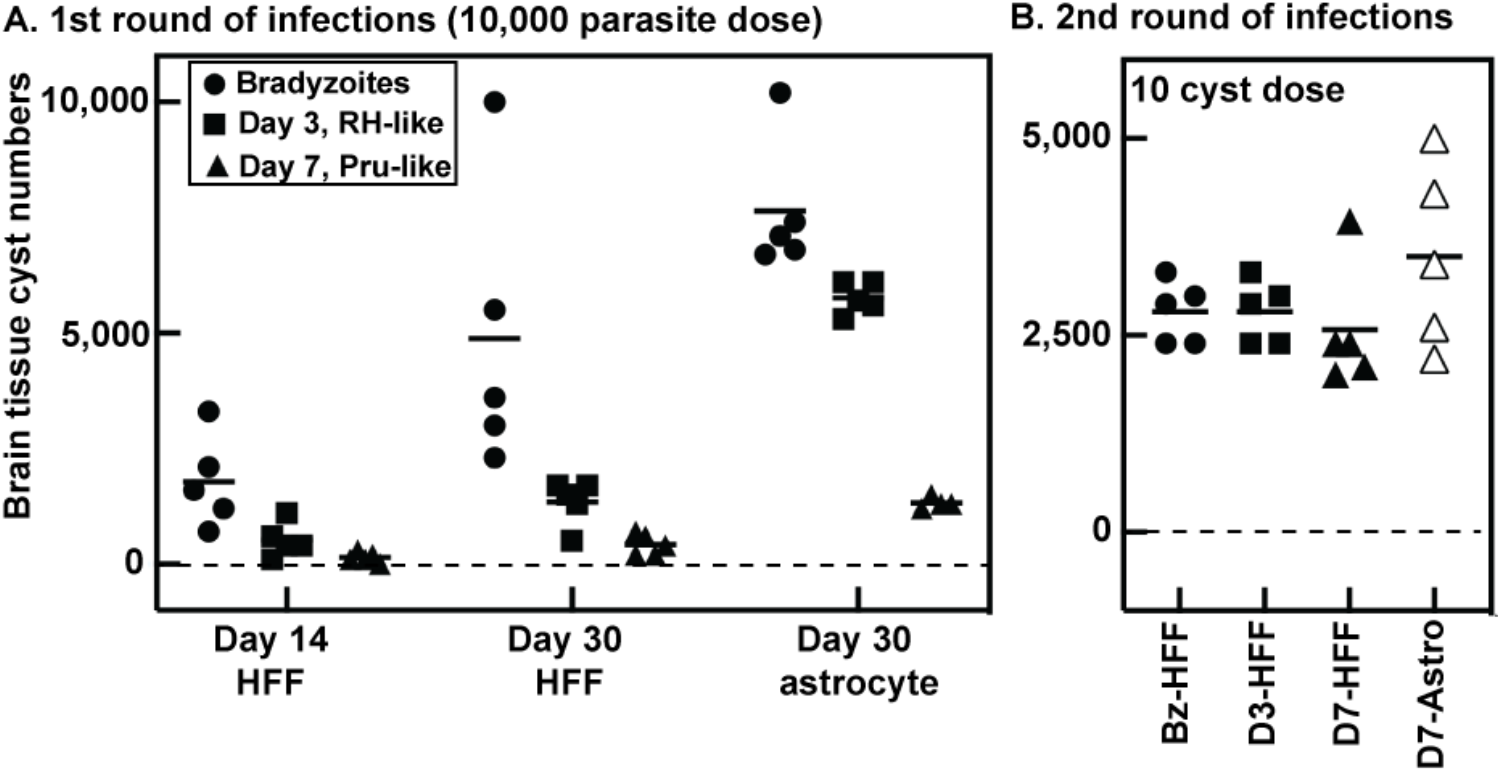
ME49EW parasites post-growth shift produce lower tissue cyst numbers in mouse brain. Native ME49EW bradyzoites were obtained from tissue cysts purified from mouse brain tissue (closed circles). Bradyzoite infection of HFF or astrocyte cells was used to obtain RH-like, Day 3 tachyzoites (closed squares). Passage of Day 5-parasites into HFF or astrocytes provided Pru-like, Day 7 tachyzoites (closed triangles). **[A.]** Groups of 5 CBA/j mice (45 total) were used to evaluate tissue cyst numbers in brain tissue at Day 14 or Day 30 PI. Infections: 10k bradyzoites/mouse inoculated i.p. with cyst counts at Day 14 and Day 30 PI (two independent cyst preparations); 10k Day 3-parasites inoculated i.p., with counts at Day 14 (only HFF-derived parasites) and Day 30 PI (HFF- or astrocyte-derived); 10k Day 7-parasites i.p., with counts Day 14 (only HFF-derived) and Day 30 PI (HFF- or astrocyte-derived). **[B.]** Tissue cysts from Day 30 PI mice infected in [A.] with bradyzoites, Day 3-parasites, or Day 7-parasites (HFF- or astrocyte-derived is indicated) were used to infect new groups of CBA/j mice (5 mice/group, 25 total); 10 cysts inoculated i.p., with tissue cyst counts at Day 30 PI.

**Figure 7.**
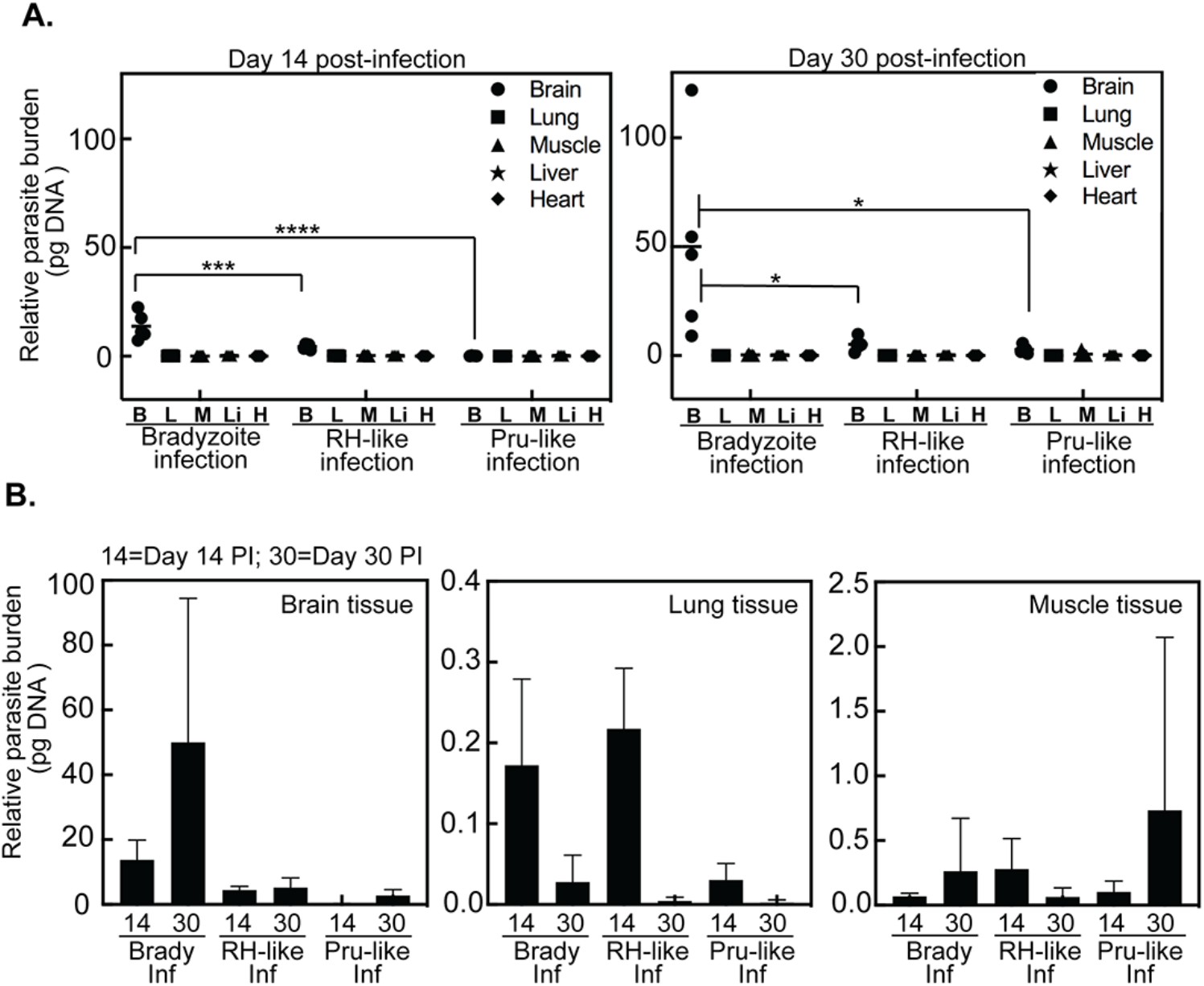
Parasite burden in mice infected with bradyzoites, RH-like and Pru-like parasites. **[A.]** CBA/j mice were inoculated i.p. with 10,000 bradyzoites, or RH-like (Day 3-populations) or Pru-like (Day 7-populations) parasites from HFF cell cultures. At day 14 and 30 PI the indicated tissues were harvested, total DNA was extracted and parasite burden determined following amplification of the B1 gene compared against a standard curve. Significance (****, p<0.0001; ***, p<0.001; *, p<0.05) is indicated. **[B.]** Relative parasite burden in three organs are compared. In brain at 14 day PI, bradyzoite vs RH-like vs Pru-like are significantly different from each other (p<0.01), while at 30 day PI only bradyzoite vs RH-like and Pru-like infected mice showed significantly different parasite burdens (p<0.05).

**Figure 8.**
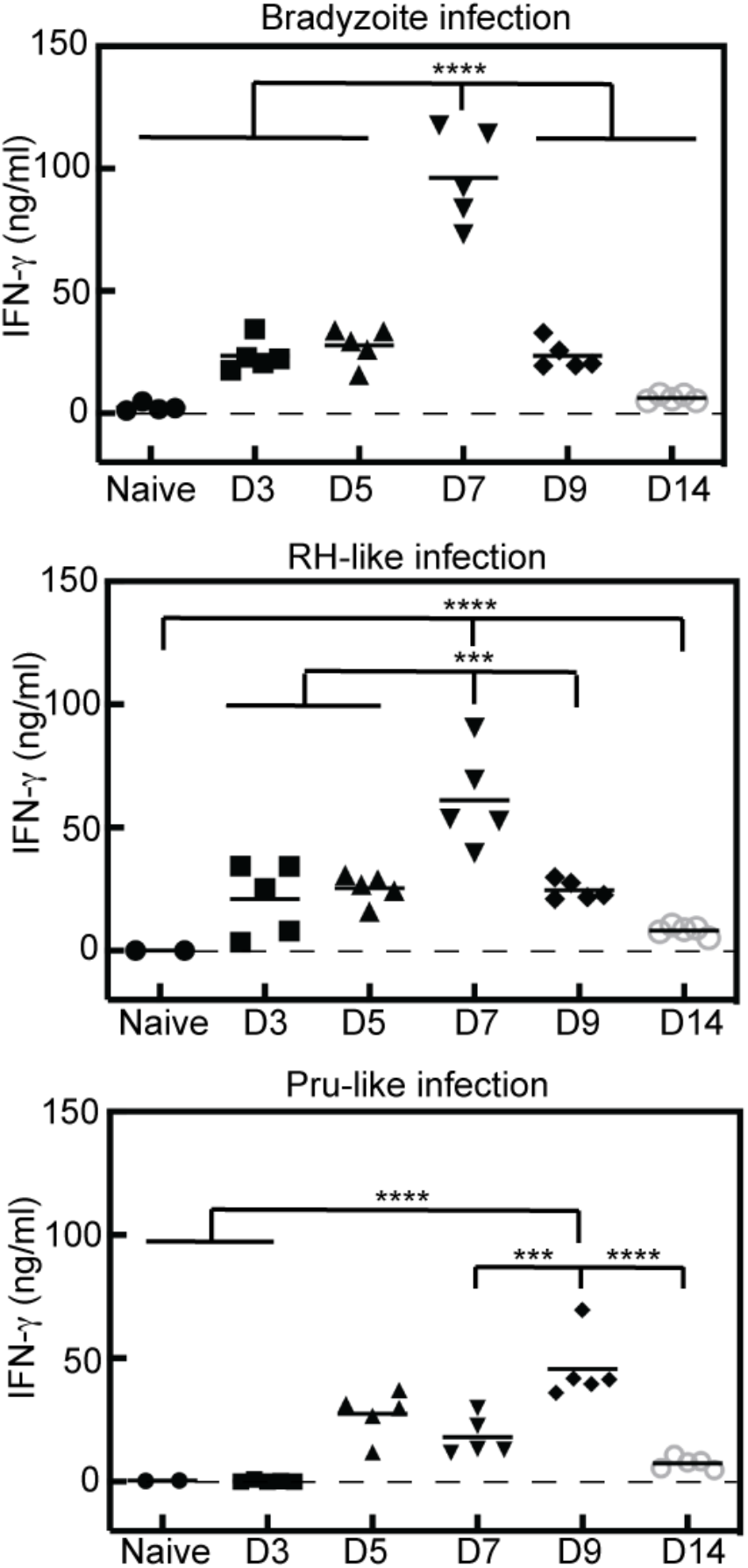
Kinetics and production of IFNγ determined by growth stage. Serum was collected from infected animals from Figure 7 at days 3, 5, 7, 9, and 14 PI with native bradyzoites, RH-like or Pru-like parasites (derived from HFF cells), and systemic levels of IFNγ were quantified. Statistical significance (*, p<0.05; **, p<0.001, *** p<0.001, **** p<0.0001) is indicated. Note also that peak IFNγ levels in the bradyzoite infection is significantly higher (p<0.02) than the RH-like parasite infection.

To determine if the reduced number of cysts in the brain are a result of poor infection and dissemination early during infection, we compared infection parameters at two different acute time points (Day 5 and 9, Fig. 9). To quantify infected cells, PECS were analyzed by flow cytometry (Fig 9). Using this method, it is possible to identify infected cells and extracellular ‘free’ parasites (29). Results revealed a distinct population of intracellular parasites (CD45+Toxo+) as well as a small but defined population of extracellular parasites (CD45-Toxo+) present in the PECS at both 5 and 9 days post bradyzoite infection. The intracellular parasite population decreased step-wise in both RH-like and Pru-like infections at 5 days post infection (dpi) (Fig. 9C,D). At 9dpi, RH-like parasites showed significantly less intracellular parasites compared to bradyzoite and Pru-like infections, which were comparable (Fig. 9D). B1 gene analysis of PECS confirmed significantly lower parasite burden in Pru-like infection compared to both bradyzoite and RH-like infections at 5dpi, suggesting a defect in the Pru-like infection (Fig. 9F). Additionally, parasite burden in the PECS significantly decreased in bradyzoite and RH-like infections between 5 and 9dpi suggesting possible dissemination and/or immune clearance (Fig. 9F). To test the presence of infection in circulation, flow cytometry was also run on blood (Fig. 9E). Extracellular parasites, previously detectable in the blood (29), could not be detected above background with any infection likely due to lower MOI. However, both bradyzoite and Pru-like infections at day 5 exhibited significantly elevated percentages of circulating infected cells compared to background. RH-like infection showed no significant increase in intracellular parasites in the blood at 5dpi indicating possible early dissemination to other tissues such as lung (Fig. 9F). By 9dpi, no intracellular parasites were found above background levels in the blood in any infection suggesting successful dissemination to other tissues. Indeed, B1 gene analysis of brain and lung tissues suggest RH-like parasites have disseminated to peripheral organs within 5 days (e.g. lung, Fig. 9F). Bradyzoite and RH-like infection demonstrated comparable B1 gene signals in brain tissue, and both were significantly elevated compared to Pru-like infection at 9dpi. In lung tissue, bradyzoite infection showed the highest B1 gene signals with RH-like and Pru-like infections showing significantly lower parasite burden. Collectively, these results indicate that bradyzoite and RH-like parasites are capable of successful infection and dissemination during acute infection, and both are able to make it to the brain by 9dpi despite enhanced immune response in the PECS. These results also suggest unsuccessful infection and dissemination capabilities of Pru-like parasites during acute infection which may explain lower cyst burdens later on during infection (see Fig. 6).

**Figure 9.**
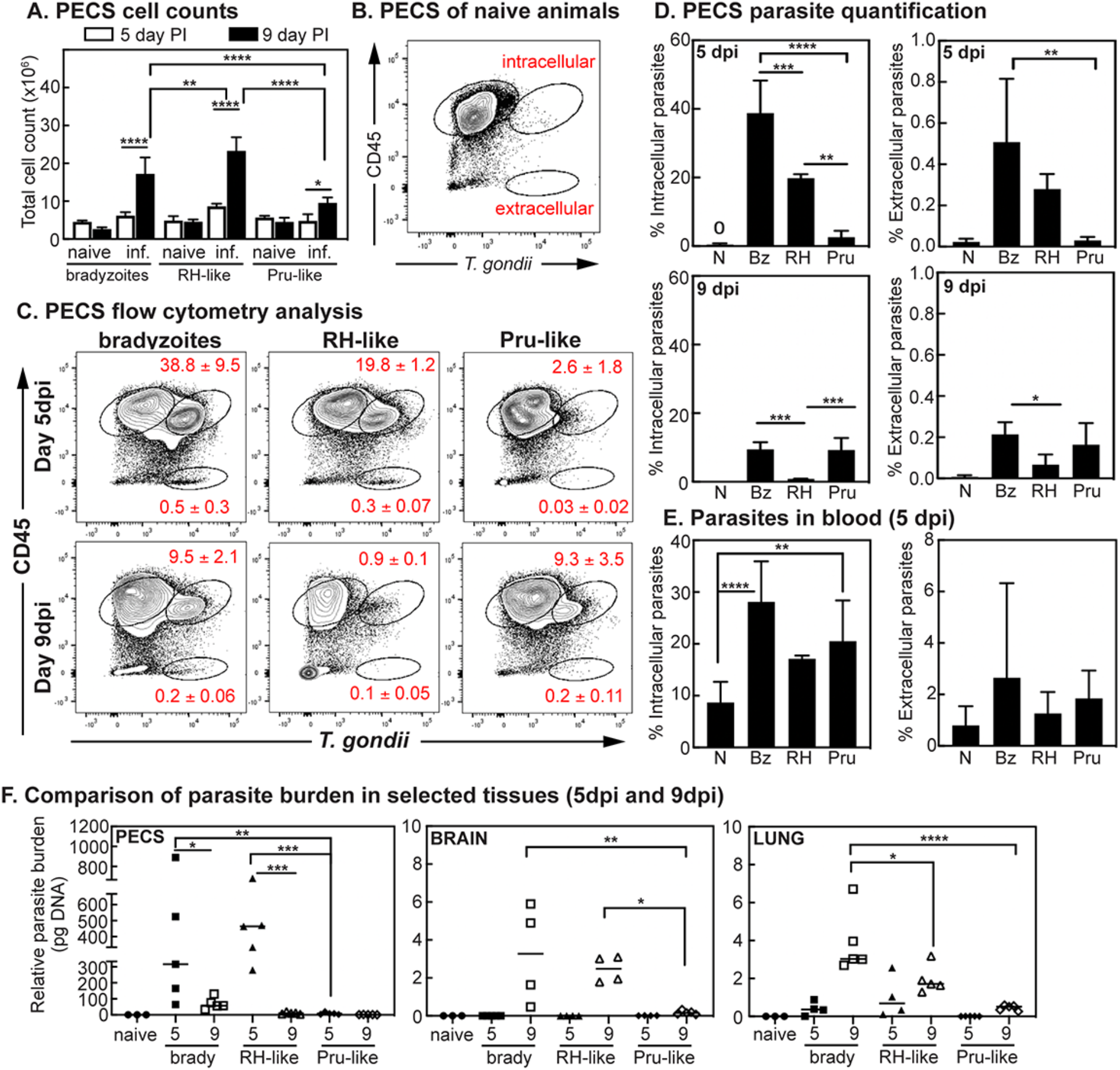
Measurement of parasite burden and dissemination during acute infection. Analysis of ME49EW acute infection results demonstrates differences in infection and dissemination capability of bradyzoites, RH-like, and Pru-like parasites. **[A.]** Cell counts from PECS of mice infected with ME49EW bradyzoites, RH-like tachyzoites, or Pru-like tachyzoites. Note the higher cell recruitment to RH-like infections. **[B.]** Flow cytometry layout of naïve PECS sample to show gating strategy. **[C.]** Flow cytometry layouts of PECS taken at 5dpi (top) and 9dpi (bottom) from mice infected with parasites from above. Numbers in red = mean ± SD for intracellular parasites (top) and extracellular parasites (bottom) (n=5/group) **[D.]** Quantification of PECS flow results. Significance determined via One-way ANOVA with multiple comparisons (p-value < 0.05). Note the greater decline of intracellular parasites from RH-like infections suggesting increased dissemination. **[E.]** Flow cytometry quantification results of blood taken at 5dpi from mice infected with parasites from above. **[F.]** Quantification of B1 gene analysis in PECS, brain, and lung at 5 and 9dpi from infected mice. Significance for all results were determined via One-way ANOVA with multiple comparisons (* p-value < 0.05, ** p<0.01, *** p<0.001, **** p<0.0001). Note that RH-like infections were detected early in lung tissue.

To further understand how bradyzoite-initiated recrudescence contributes to tissue cyst formation, we infected CBA/j mice with native ME49EW bradyzoites and then blocked parasite growth by adding sulfamerazine to drinking water at various times (Fig. 10, top diagram). Groups of five mice were infected with 10,000 bradyzoites and subjected to sulfamerazine treatment for a period of ten days starting three days PI and continuing until day 12 PI. At 30 days PI, the brain cyst numbers were determined (Fig. 10). The addition of sulfamerazine to drinking water at Day 3 PI effectively reduced tissue cyst numbers by >400 fold over no drug controls, while addition at Day 9 PI reduced average cyst numbers 13 fold. Treatment started at 12 days PI had no significant effect on average cyst numbers in 30 day infected mice. These results were consistent with the finding that bradyzoites and RH-like populations that formed early in bradyzoite recrudescence have the greatest capacity to produce brain tissue cysts in CBA/j mice (Fig. 6). Importantly, these results also demonstrate that the complete series of steps in this developmental pathway, which can not be duplicated by lab-adapted strains, are critical to achieve high cyst numbers in mouse brain tissue.

**Figure 10.**
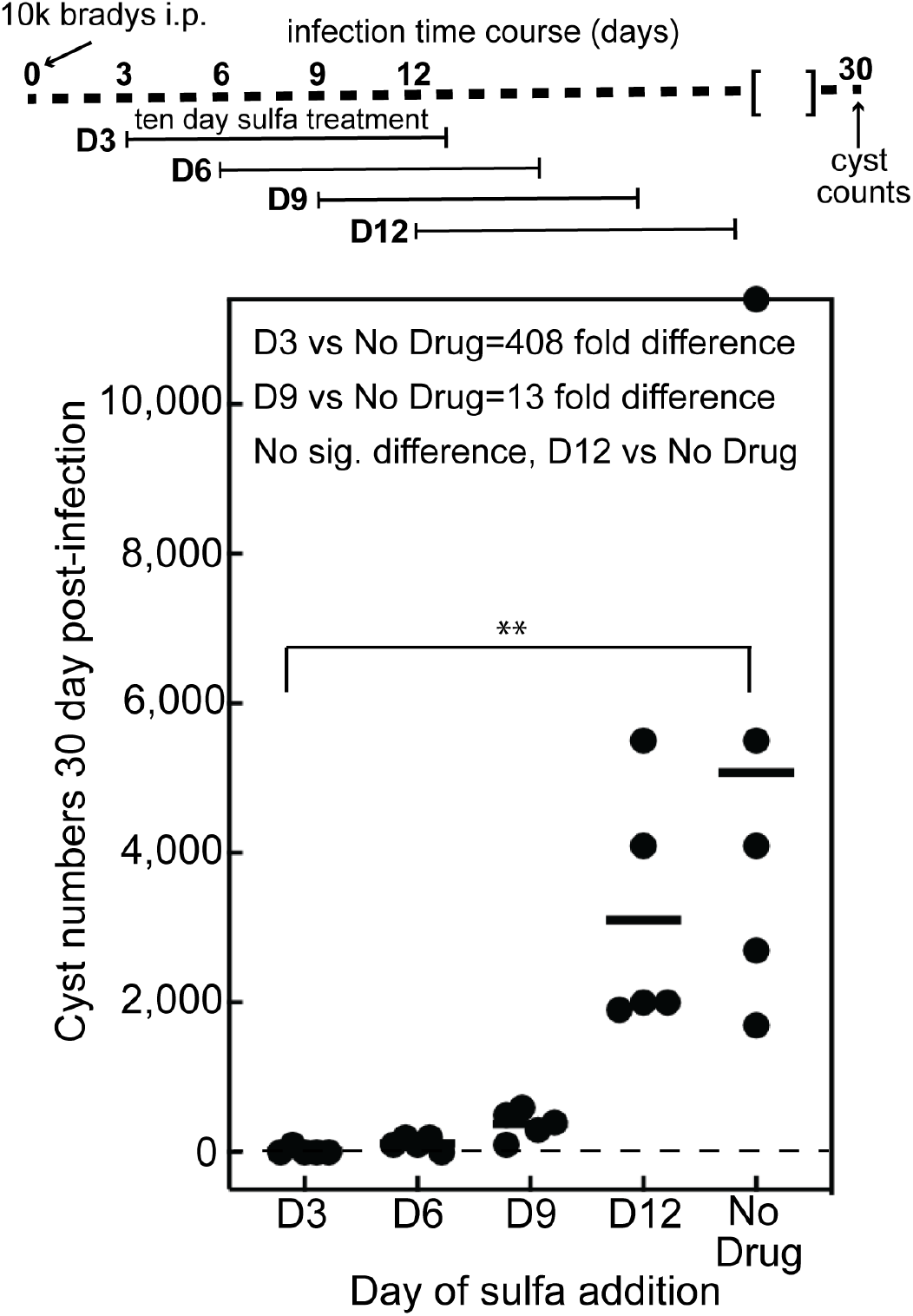
Early growth expansion of ME49EW parasites is critical for tissue cyst formation in mice. Five groups of 5-CBA/j mice (25 total) were infected with 10k purified native ME49EW bradyzoites. The times of sulfamerazine (sulfa) addition and duration of treatment are indicated above the graph. Tissue cyst numbers per mouse brain were determined at 30 days PI. Statistical significance (**, p<0.001) is indicated. Note: The fold decrease in cyst number was >400 lower when the drug was added 3 days PI as compared to the no drug control and 13 fold lower when drug was added 9 days post-infection.

## DISCUSSION

Why some *Toxoplasma* strains produce thousands of tissue cysts in mouse brain, while other strains produce a couple of dozen is not known but has been a bottle neck in advancing our understanding of cyst biology. The native ME49EW and lab-adapted PruQ parasites studied here are largely equivalent in the alkaline-stressed in vitro model; both strains slow growth in pH8.2 media, and robustly assemble cyst walls in 48-72h cell cultures. However, where it really counts in *Toxoplasma* infections these strains differ by two orders of magnitude in the ability to produce tissue cysts in mice (Fig. 1). It is well known that repeated passage in HFF cells exacerbates the loss of developmental competency (reviewed in refs. 11, 30), therefore, even if developmental competency is only partially affected after the initial HFF-adaptation, it is likely to be fully lost over time in HFF cell cultures. The dilemma for the field is to decide when HFF-adapted strains are a suitable choice for developmental study and when they are not. It is worth noting that the biological changes associated with developmental losses provide clues to the possible underlying mechanisms : 1) lab-adapted parasites do not appear able to deeply growth arrest like native bradyzoites and instead they become locked in a replicative cell cycle that becomes faster with sequential passages, therefore, G0/G1 mechanisms must be involved. 2) As we showed here, lab-adapted parasites gain the ability to survive outside the host cell for longer periods, which others have reported is associated with stress resistance (26). Finally, 3) the inability of lab-adapted strains to fully develop into mature bradyzoites is significantly correlated with poor cyst production in mice. We replicated many of these changes in generating the lab-adapted ME49EW1 strain, although six separate platings of in vivo ME49EW bradyzoites in HFF cells were required for this strain to spontaneously emerge. These results suggested the worrisome idea that ME49EW1 parasites arose due to genetic mutation; an idea that was reinforced by a small mutagenesis experiment (Alvarez and White, personal communication). Using mutagenesis conditions that previously generated temperature sensitive *Toxoplasma* mutants (31), we exposed ME49EW bradyzoite-infected HFF cultures to ethyl nitrosourea, which led to the emergence of multiple HFF-adapted parasite lines from a single bradyzoite plating (2 million bradyzoites). Taken together, these results indicate there is an underlying genetic basis that causes developmental losses to become permanent in many strains like PruQ and ME49EW1, whereas, the temporary loss of developmental competency we observed for the ME49EW “Pru-like” recrudescent population is likely due to epigenetic changes that are fully reversible. While we do not understand the molecular basis for the permanent loss of developmental competency of lab strains, we demonstrated molecular markers that distinguish the developmental endpoints of in vitro versus in vivo models can be identified and these new markers could be used to eliminate strains that fail to properly develop.

The relatively few studies of bradyzoite recrudescence in mice and humans have uncovered evidence that bradyzoite recrudescence is nonsynchronous (8, 9) resulting in actively replicating tachyzoites and all stages of bradyzoite re-development from immature to mature cysts being present in the same lesion (8). Here we have introduced an innovative ex vivo model of bradyzoite recrudescence that captures all these features with greater cell biologic and temporal resolution (see Fig. 11 model). Studies spanning 30 years support the concept that the cell cycle end point of tachyzoite to bradyzoite development is dormancy (10, 20, 32), which is substantiated by our study. Our results confirm that mature ME49EW bradyzoites (>30 days PI) are deeply growth arrested; they have 1N genomic contents and lack mitotic and budding forms, have downregulated many growth-related gene transcripts, and are very slow to resume replication after invasion. We do not understand the regulatory basis of bradyzoite dormancy, however, we believe these mechanisms are important for limiting recrudescence as well as cyst size and number. Remarkably once awakened, development initiated by ME49EW bradyzoites follows a similar pathway that we previously described for sporozoite infections (24, 33). ME49EW bradyzoites sequentially develop into two types of tachyzoites; a fast-growing tachyzoite (RH-like) that switches in days to a slow-growing tachyzoite (Pru-like). The rapid differentiation of ME49EW bradyzoites into a fast-replicating tachyzoite tracked with the same timeframe in both fibroblasts and astrocytes as did the shift to slower growth indicating these growth changes are host cell-independent. The new ex vivo model of bradyzoite recrudescence also confirmed that an alternative bradyzoite-bradyzoite replication pathway exists in the intermediate life cycle that had been inferred by other research (34, 35). Our results revealed astrocytes are permissive for bradyzoite replication while fibroblasts are resistant. The basis for this cell-specificity is unknown. ME49EW bradyzoites occupy single parasite SRS9+SAG1-vacuoles for ∼24h and it takes ∼5 days to fully transition to a near 100% SAG1+ tachyzoite culture in fibroblasts. In contrast, bradyzoites that invade primary astrocytes exhibit two forms of replication simultaneously that follow the 24h reawakening period: bradyzoite to tachyzoite conversion and replication (∼85% vacuoles) and bradyzoite to bradyzoite replication (∼15% vacuoles). It is often assumed that *Toxoplasma* developmental progression primarily occurs in response to immune stress. The results here demonstrate that the course of growth and developmental changes in the intermediate life cycle are the result of parasite-encoded mechanisms responding to signals from the host intracellular environment and are independent of classical immune-mediators.

**Figure 11.**
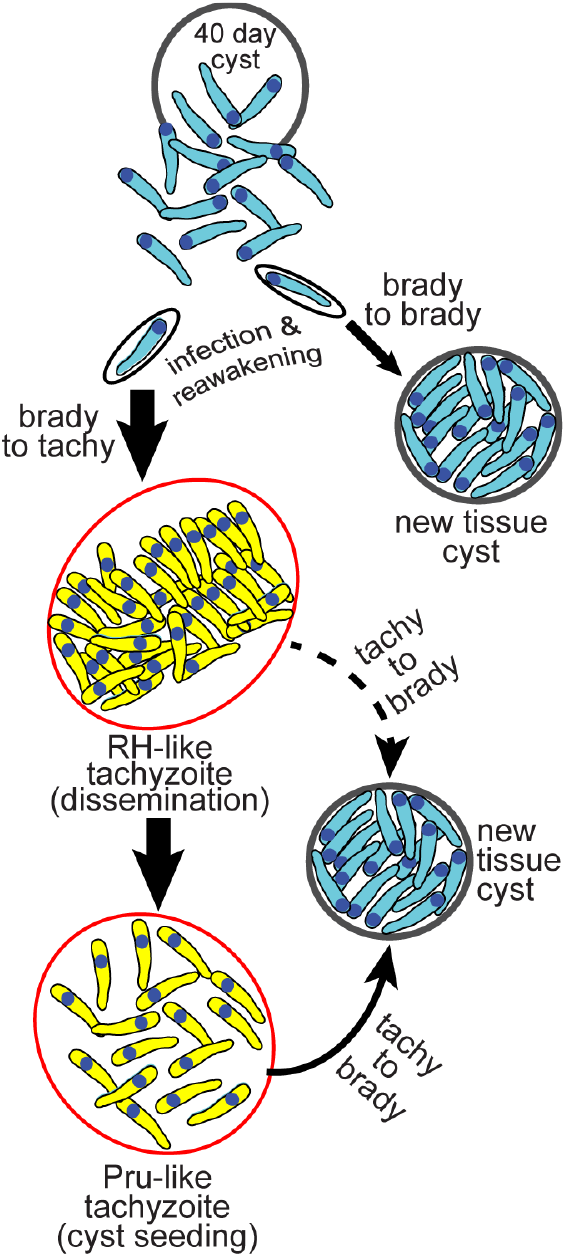
Model of *T. gondii* tissue cyst recrudescence. Dormant brady-zoites from mature tissue cysts are capable of initiating two distinct developmental pathways.

For close to 100 years studies of the *Toxoplasma* tachyzoite have been conducted using in vitro cultured fibroblasts, indeed the ease of this culture system facilitated the study of *Toxoplasma* tachyzoites as a model for apicomplexan biology. In contrast, our new ex vivo model of bradyzoite recrudescence captures two distinct pathways, one host cell dependent and the other independent. Host cell dependency has been a feature of *Toxoplasma* since the discovery of spontaneous cyst development occurring in neurons and myocytes (36). Yet the heterogeneity achieved in astrocyte cultures of tachyzoite and bradyzoite replication is new and suggests a major portion of the life cycle that has been overlooked. *Toxoplasma* encounters many cell types during its lifecycle some of which induce cyst production and some that facilitate dissemination. Recrudescence most commonly occurs at three different sites i) in the gut following ingestion of tissue cysts where parasites encounter gut epithelial cells followed by innate immune cells ii) in peripheral organs such as the muscle and liver and iii) in the brain/CNS containing neurons, astrocytes, microglia and, during infection, peripheral immune cells. Determining what characteristics of the astrocyte support bradyzoite-bradyzoite replication and whether such characteristics exist in other cell types the parasite encounters will have important implications for managing the amplification of bradyzoites and the development of cysts. Of note we have documented bradyzoite replication in astrocytes in a culture dish devoid of IFNγ. This cytokine is critical for protection against toxoplasmic encephalitis (TE) in part by limiting parasite replication in astrocytes (37, 38) indeed in the absence of IFNγ signaling, cyst formation is observed in astrocytes (22, 38). Our ex vivo model by no means solves the question of what prompts a cyst to reactivate, however, in the context of TE (absence of IFNγ) and during initial infection the developmental pathway uncovered by this ex vivo model suggests the parasite is pre-programmed to undertake two important functions: 1. Amplify parasite burden to maximize dissemination and 2. Seed tissues with cysts that will continue transmission. These functions occur not sequentially driven by immune responses but are pre-programmed hence cyst formation even in the absence of IFNγ. Recrudescence is therefore a point in the lifecycle in which *Toxoplasma* has the potential to go from a homogeneous to a heterogeneous population composed of both replicating bradyzoites and tachzyoites in the same environment. We propose that this heterogeneity is functional. Bradyzoite to bradyzoite replication allows for the growth and maintenance of the cyst population either in neurons resulting in larger cysts and ‘twin cysts’ or via astrocytes as an energy-rich vessel facilitating optimal brady-brady replication and local cyst dissemination. Although significant progress has been made using genetically modified lab-strains to understand transcriptional regulation, cyst location, number and kinetics in vivo, in the absence of the complete recrudescence pathway (quickly lost in vitro), these studies are likely testing the initial seeding of cysts in the brain. Our results show that the complete bradyzoite to tachyzoite recrudescence pathway that unfolds over one week following purified bradyzoite infections is required for robust expansion, dissemination, and establishing tissue cysts in brain tissue. This window of opportunity is consistent with a recent study showing maximum tissue cyst formation in pig muscle is reached seven days after oral infection with tissue cysts from mice tissues (39). Infections with the 3 growth forms of the excysted ME49EW bradyzoite paints a picture of recrudescence following tissue cyst infection. Efficient bradyzoite infection of cells followed by a rapidly dividing phase after 24h leading to almost 40% of cells infected by day 5 and a measurable number of extracellular parasites disseminating in the blood and migration to tissues with the lung showing early signs of infection. For the next four days ‘RH-like’ parasites are seeding tissue especially the brain at which point they will encounter neurons encouraging cyst formation. By this time high systemic IFNγ activates host cells to limit parasite replication and clear this lytic form of infection. The pre-programmed switch to a slower growing tachyzoite stage by day 7-9 enhances both the immune clearance of the acute infection and promotes conversion to the bradyzoite stage. This timing is supported by our data showing that drug treatment after day 9 is ineffective at limiting cyst burden confirming the slowing of parasite growth effectively ends dissemination. If the complete pathway is not intact and the fast growing ‘RH-like’ phase is skipped, as is the case when we infect with ‘Pru-like’ parasites, then cysts in the brain are >50-fold lower than infection with purified bradyzoites. The viability of these cysts is intact demonstrating that the parasites are cyst-competent. Instead it is the slow growth and poor dissemination of this slow-growing stage of the recrudescent pathway supported by fewer infected cells and a delay and reduction in the magnitude of the host response that likely accounts for low cyst numbers. We postulate that many of the lab-adapted strains that remain cyst-competent are stuck in this ‘Pru-like’ phase of growth. In contrast, purified bradyzoites that remain single undivided parasites for 24h result in a greater cyst burden in the brain over the ‘RH-like’ rapidly dividing parasite. Of note, the bradyzoite infections exceed ‘RH-like’ in almost every parameter except the number of cells recruited to the site of infection. This could suggest a ‘stealth-like’ component of the excysted bradyzoite stage that would be missed in the rapidly dividing and lytic (immune-amplifying) ‘RH-like’ infections or an enhanced capacity to manipulate host cell migration thereby increasing dissemination as has been documented for tachyzoites (40).

In summary, the dormant bradyzoite in ME49EW tissue cysts possesses remarkable developmental flexibility and ability to tailor the parasite-encoded developmental program to the type of host cell infected, which is fully captured in a new ex vivo astrocyte recrudescence model. In the right host cell, ME49EW parasites can chose to replicate as bradyzoites or fully convert to the tachyzoite stage in order to disseminate. Native bradyzoites, like the sporozoite stage, develop into the tachyzoite stage using a two-step pathway that permits rapid biomass expansion but is also self-limiting. These versatile tachyzoites also have the ability to re-develop into the dormant bradyzoite at any time within a host cell environment that is permissive for tissue cyst development. Adapting native strains to HFF cell culture disrupts these pathways and may cause transcriptional confusion as recent single parasite RNA-seq studies have suggested (41).

## MATERIALS AND METHODS

### Parasite strains and host cells

All laboratory-adapted strains were maintained in human foreskin fibroblasts (HFF) cells in high glucose DMEM with sodium bicarbonate (Sigma) supplemented with 5% heat-inactivated fetal bovine serum (FBS) and 1% penicillin-streptomycin in humidified incubators at 37°C with 5% CO2 and ambient oxygen (21%). The Type II ME49EW strain (kindly provided by Emma Wilson, U. California Riverside) was exclusively maintained in mice as described in Figure S1 and below. The strain ME49EW1 was obtained by forced adaptation of the parent ME49EW strain to grow well in HFF cells under standard culture conditions used to grow HFF-adapted strains and then cloned by limiting dilution. Several other HFF-adapted laboratory strains were used in this study: RH (Type I RH Δ*hxgprt*)(42), GT1 (low passage Type 1 strain)(43), PruQ (Type II Prugniaud Δ*ku80*, Δ*hxgprt::*LDH2-GFP)(13), and ME49-Tir1 (Type II ME49-Tir1-3 Flag)(12).

#### Astrocyte purification

Neonatal C57Bl/6 mice born at 0-3 days were sacrificed via decapitation and brains (without cerebellum) dissected and placed in cold wash media (DMEM and 2% fetal bovine serum). Brains were pooled and homogenized through a sterile 40 micrometer cell strainer (Corning) using a 3 ml syringe plunger. Homogenate was washed twice with cold wash media and spun at 2,000 rpm for 10 minutes at 4°C. Homogenate was resuspended with 6 ml per brain of prewarmed complete media (DMEM [Corning]; 10% FBS; 1% Nonessential Amino Acids Mixture 100x, [Biowhittaker Reagents Lonza],1% GlutaMAX supplement 100x [Thermo Fisher Scientific], 1% penicillin-streptomycin, [Genclone], 1% HEPES Buffer solution 1M [Thermo Fisher Scientific]. Resuspended cells were plated in non-vented T-25 cm^2^ flasks with loose caps at 37°C, 5% CO2. At days 3, 5, and 7 astrocyte media was replaced. To remove contaminating less adherent microglia and oligodendrocytes, astrocytes were shaken on day 8-10 at 260 rpm, 37°C for 1.5 hours. Media was replaced and cultures were shaken for an additional 24h at 100 rpm, 37°C. Astrocytes were lifted with 3ml of 0.25% Trypsin EDTA, counted, and plated at a density of 1×10^6^ cells per T-75cm^2^ vented flask until confluent.

### Animal experiments

All animal research was conducted in accordance to the Animal Welfare Act. C57Bl/6, CBA/J, and SWR/J #689 mice obtained from Jackson Laboratory (Jackson ImmunoResearch Laboratories, Inc., West Grove, PA, USA) were bred and maintained in a pathogen free vivarium under the Institutional Animal Care and Use Committee (IACUC) protocols established at the University of California, Riverside and the University of South Florida, Tampa.

The ME49EW strain was continually passed through 5 to 6 week old female mice by first infecting resistant strain SWR/J (Swiss Webster) with 10 cysts in 200μl of cortex homogenate by intraperitoneal (i.p.) injection (see Fig. S1 for details). Tissue cysts in brain homogenates from infected SWR/J mice were then used to infect CBA/J, sensitive mice (10 cysts in 200μl diluted brain homogenates, i.p.).

#### Brain homogenate preparation

At 14-60 days PI as indicated, infected SWR/J or CBA/J mice were euthanized, brain cortices dissected and placed in 3ml of cold PBS (cerebellum was discarded as it contained <5% of total brain tissue cysts). Tissue homogenization and tissue cyst yields were significantly improved (tissue cyst numbers >50% higher) by storing infected brain cortices overnight at 4°C in PBS. In direct comparisons, we determined tissue cysts held in brain tissues for up to 24h in cold PBS were viable, fully infectious, and yielded active excysted bradyzoites. After the 4°C overnight storage in 3ml of 1xPBS, cortexes were more easily serially homogenized by 3-4 serial passes through 18, 20 and finally, 22 gauge (blunt) sterile needles. Tissue cyst quality and enumeration was determined by a full examination of 30μl (x 3) of homogenate spread on a microscope slide. All tissue cyst mouse inocula used in this study were diluted to 10 cysts in 200μl in 1xPBS. In addition, the quantification of brain tissue cyst counts in mice infected with various parasite sources followed these protocols and were determined in triplicate.

#### Tissue cyst purification and bradyzoite excystation

Cysts were purified from mouse brain cortices infected with the indicated parasite numbers/cysts for 30-50 days. To purify tissue cysts, brain cortices were harvested and homogenates prepared as above (one cortex per 15 ml conical tube as combining cortices reduces cyst yields). Following cyst counts as above, homogenate volumes were brought up to 11ml with PBS, mixed by inversion, and then centrifuged at 500xg for 15 minutes. The supernatant was removed by aspiration and the homogenate pellet resuspended in 2.5ml of percoll-90 (90% percoll, 10% 1xPBS) followed by the addition of PBS to a final volume of 7.5ml (final percoll=30%). The tube was fully mixed by several inversions and then centrifuged at 1000xg for 15 minutes without a brake. Tissue cysts pelleted in this gradient protocol, while debris collects at the top of the gradient, therefore, the supernatant was carefully removed by aspiration. If multiple cortices are used, the pellets from the individual percoll gradients that contain tissue cysts were combined and the total volume raised to 10ml of PBS, followed by centrifugation at 30xg for 10 min and then removal of the supernatant by careful aspiration. A low speed 30g spin pellets medium to large tissue cysts (majority of bradyzoites) while >80% of red blood cells are retained in the supernatant. To obtain free bradyzoites, the tissue cysts were resuspended in excystation solution (5mg/ml pepsin, 100mg/ml NaCl, 0.14N HCl), incubated for 90 seconds and then 10 ml of culture media containing 5% fetal bovine serum added. Bradyzoites counts were obtained using a hemocytometer. A more detailed protocol with tips for successfully producing and purifying tissue cysts and free bradyzoites is available upon request.

#### Infections of mice with laboratory strains and recrudescing populations

ME49EW bradyzoites and parasites at various times during recrudescence in HFF or astrocyte cells, as well as laboratory strains grown in HFF cells, were purified from host cell monolayers by standard scrape, needle pass, and filter methods and diluted to 50,000 parasites/ml in PBS. We have determined experimentally that filtration of needle-passed parasite cultures through 3 micron filters removes >95% of trypan blue positive (dead) parasites. Groups of CBA/J mice as indicated were infected i.p. with 0.2ml of the appropriate inocula. Mouse infections were used to determine parasite burden, cytokine expression and tissue cyst formation.

#### Serum Cytokine ELISA

Serum was harvested from mice via tail vein bleeding on days 3, 5, 7, 9, & 14. Blood was centrifuged at 16,000 RPM for 20min. Serum was aliquoted and stored at 80°C. IFNγ capture antibody (ebioscience: 14-7313-81) was incubated overnight at 4°C [1:1000]. Recombinant IFNγ (ebioscience, cat#: 14-8311-63) was used to create the standard. Standard and serum samples were incubated at 37°C for 2h. Biotin conjugated anti-mouse IFNγ (clone:R4-6A2; ebioscience13-7312-85) was incubated at 37C for 1h. Peroxidase conjugated streptavidin (Jackson immune research 016-030-084) was incubated for 30min at 37°C. TMB substrate (Thermo, N301) is used for the colorimetric reaction, and an equal part of 0.16M sulfuric acid was used to stop the reaction. Standard curve and analysis were done in GraphPad Prism (GraphPad Software).

#### Sulfamerazine treatment

CBA/j (25 mice total) mice were each inoculated with 10,000 excysted ME49EW bradyzoites by i.p. injection. At various times following infection (see Fig. 9 for details) sulfamerazine (444 mg/liter) (44) was added to the drinking water of selected groups of mice. At 30 days PI, mice were euthanized, brain cortices removed and tissue cysts in homogenates enumerated in triplicate.

### Ex vivo bradyzoite recrudescence experiments

Preliminary investigations determined that native ME49EW parasites grew better in HFF cells under lower oxygen levels. Therefore, throughout this study we cultured ME49EW parasites in HFF or primary astrocyte monolayers that were placed in a hypoxic chamber and grown in the same media with the exception that astrocytes received a 1% GlutaMAX supplement (see Fig. S1 for further media details). Confluent HFF or neonatal astrocyte monolayers were inoculated at various times with excysted bradyzoites, lab strains or recrudescing populations at a MOI ∼0.5. ME49EW parasites were passaged into new monolayers of either host cell type using the schedules shown in Figure S2A.

#### Growth rate doublings

Post-excysted bradyzoites and Day 3- and Day 5-recrudesing populations were inoculated onto 6 well coverslips of HFF or astrocyte cells and at various times were fixed and stained using an anti-toxoplasma antibody (abcam, ab138698). Vacuole sizes at each timepoint were determined by counting 50 random vacuoles in triplicate on a Zeiss Axiovert fluorescent microscope. For each coverslip, the average number of parasite doublings per day was determined from the raw vacuole sizes (vac of 2=1 doubling, vac of 4=2 doublings, vac of 8=3 doublings etc.) divided by the total days in each incubation period.

#### Visualization of parasite growth and development

ME49EW bradyzoites and various recrudescing populations were inoculated onto 6 well glass coverslips of HFF or astrocyte cells at 0.5 MOI. All coverslips were cultured under low oxygen conditions as described in Figure S1. Infected coverslips were fixed with 4% PFA for 10min. Cells were permeabilized with either 100% acetone or 0.25% Triton-X for 10min. Cells were blocked with either 5% donkey serum or 1% BSA for 30min, followed by a 1h incubation of the following primary antibodies diluted in blocking buffer: biotinylated-dolichos biflorus agglutinin (DBA) [1:500] (Vector Laboratories), rabbit-anti SRS9 [1:1000] and mouse-anti SAG1 [1:1000](kindly provided by John Boothroyd, Standford U.), rabbit-anti-Toxo [1:500] (Abcam), mouse-anti-CST-1 [1:1000](kindly provided by Louis Weiss, Albert Einstein COM), mouse-anti-IMC1[1:1000](kindly provided by Gary Ward, U. Vermont), rabbit-anti-centrin [1:500](used 2 hour primary incubation). Secondary antibody master mix were incubated for 1h: Goat-anti-rabbit-AF568 [1:1000], Goat-anti-mouse-AF488 [1:1000], streptavidin-AF647 [1:1000], DAPI (1mg/ml). All incubations were done at room temperature, and all washes used 1XPBS.

#### Extracellular viability

Day 2-recrudesing ME49EW parasites (bradyzoite infection of HFF or astrocytes) and lab-adapted strains, ME49EW1, RH and GT1 parasites were purified and resuspended in 37°C growth media (no host cells). The host-free parasites (10,000/cover slip) were used to immediately inoculate cover slips of the host cell type they were obtained from, and then again, at 6, 12, and 24h (only 0, 6, and 12h are shown). Parasites were incubated at 37°C throughout the experiment and ME49EW parasites were cultured in hypoxic chambers, while lab-adapted strains were cultured under standard ambient oxygen conditions. Infected coverslips were left to grow undisturbed for ∼4 days prior to being fixed and IFA analysis performed using anti-toxo primary antibody. Microscopic quantification of parasite plaques was determined by counting the number of vacuoles per field for a total of 10 fields (in triplicate). Percent survival was determined in comparison to the time 0 infections.

### Cell Prep and Flow Cytometry

#### Cell Preparation

PECS and blood were collected from naïve and infected mice at each time point using injection and reuptake of 1xPBS using a 22G needle and 5ml syringe (for PECS). Volume collected was recorded. Blood was collected by cutting right atrium of heart with scissors and collecting approximately 500µl. Blood was subjected to 2 rounds of ACK Lysis Buffer (500µl) and pipetted through filter cap to minimize debris. PECS and blood cells were counted using an automated cell counter and 1.0×10^6^ live cells were taken for flow cytometry staining.

#### Flow Cytometry Staining

During cell prep and flow cytometry staining, all spins were conducted at 3000rpm for 5min at 4°C to spin down extracellular parasites as well as infected cells. All cells were incubated in FC Block (BD Pharmingen: 553142) to minimize non-specific antibody binding, surface stained with CD45 antibody conjugated to PE (eBioscience: Clone 30-F11) [1:100] to identify immune cells, fixed with 4% PFA (15 min at room temperature), and permeabilized via 0.3% Saponin spin for intracellular staining. Cells were then intracellularly stained with primary Rabbit anti-Toxo antibody (abcam: ab138698) [1:200] followed by Goat anti-rabbit AlexaFluor488 secondary stain (eBioscience) [1:500] to identify both intracellular and extracellular parasites. All intracellular stain steps were completed in 0.3% Saponin to maintain permeabilization. In between antibody stains, cells were spun down to discard any unbound antibody. Upon completion of intracellular staining, cells were spun in FACS buffer to close membrane, and cells were resuspended in 300µl FACS buffer for flow cytometry. Flow cytometry was run on BD FACS Canto II, and analysis was performed using FlowJo software.

### Parasite burden

Organ tissues (brain, lung, muscle, liver, heart) and peritoneal exudate cell suspension (PECS) were harvested from mice and stored at -80°C prior to DNA purification. DNA was purified using a DNeasy minikit (Qiagen) according to manufacturer protocols. The qPCR reaction volume used 600ng of DNA from each organ, 0.375μM B1 gene primers (Forward:5′-TCCCCTCTGCTGGCGAAAAGT-3′; Reverse: 5′-AGCGTTCGTGGTCAACTATCG-3′), and 2x Luna universal qPCR master mix (New England Biolabs) in a 25μl total volume. The amplification program was: 3 min at 95°C, 40 cycles of 10s at 95°C for denaturation, 30s at 60°C for annealing, followed by 5s at 65°C and 5s at 95°C for extension. The realtime data was collected on a CFX96 Real-Time system, C1000 Touch thermal cycler using Bio-Rad CFX maestro software. A standard curve was generated based on serial dilution of purified ME49 genomic DNA. All results were expressed as the pg of parasite genomic DNA calculated from an experimentally generated standard curve generated in each PCR run.

### Gene expression

#### RNA purification

Total RNA from ME49EW cysts, ME49EW1 tachyzoites and Day 1-, 2-, or 7-recrudescent populations as well as PruQ tachyzoites and alkaline-induced PruQ bradyzoites (pH8.2 media for 72 h) were purified using the RNeasay kit (Qiagen) according to manufacturer protocols. Two biological replicates were prepared for each RNA source and RNA quality was assessed using an Agilent 2100 Bioanalyzer (Waldbronn, Germany).

#### RNAseq analyses

RNA samples were processed for RNA-sequencing using the NuGen Universal RNA-Seq System (NuGEN Technologies, San Carlos, CA). Briefly, 100 ng of RNA was used to generate cDNA and a strand-specific library following the manufacturer’s protocol. In order to deplete contaminating abundant human and *T. gondii* rRNA sequences, a cocktail of NuGEN human AnyDeplete rRNA-targeting probes and a custom panel of 196 *T. gondii* AnyDeplete probes were used in order to carry out insert-dependent adaptor cleavage, as described in the manufacturer’s protocol. The final cDNA libraries were evaluated on the BioAnalyzer and quantified using the Kapa Library Quantification Kit (Roche Sequencing, Pleasanton, CA) by Molecular Genomics Core Facility (Moffitt Cancer Center, FL). The libraries were then sequenced on the Illumina NextSeq 500 sequencer with 75-base paired-end runs in order to generate at least 10 million read-pairs per sample. Reads were filtered by Sanger quality score using FASTQ Groomer v. 1.0.4 and paired end reads were aligned against the genomes of *T. gondii* (TGME49 version 46; ToxoDB.org) references uploaded into Galaxy using HISAT2. The DESeq2 module was used for normalization, differential gene expression and statistical analysis of uniquely mapped paired-end reads using the default parameters.

#### mRNA qPCR

Recrudescing populations were used to infect HFF cells (in duplicate) with 1.5-2 million parasites (MOI 1:1), parasites grown for 24-48h, and then monolayers were washed in ice cold 1xPBS, scraped on ice in 1xPBS and pelleted at 3800rpm for 15 min at 4°C. The pellet was resuspended in 350μl of RTL buffer and total RNA was purified by manufacturer protocols (RNAeasy kit, Qiagen) RNA was converted to cDNA and purified from each sample by standard methods. The qPCR reaction was: 10μl Luna Universal qPCR master mix (New England bio labs), 0.5μl Primer (of a 10μM stock), 20ng of tachyzoite cDNA or 10ng of cyst cDNA in a 20μl total volume. The amplification program used was 3 min at 95°C, 40 cycles of 10s at 95°C for denaturation and 30s at 60°C for annealing followed by 5s at 65°C and 5s at 95°C for extension finally held at 4°C. See Dataset S1 for all primer designs.

Calculations: Sample 2^-ΔCt^=2^-(Ct target gene-Ct of ADF control)^

Relative fold change=2^-ΔCt^ sample target gene/2^-ΔCt^ Pru-7 value for the same target gene

### Cell cycle analyses

#### DNA Content

Excysted bradyzoites and freshly lysed and purified RH parasites were pelleted at 2000 rpm for 15 min at 4°C and resuspended in 300μl of cold 70% ethanol in PBS added dropwise. Samples were held at -20°C at least 24h prior to staining. Samples were pelleted by centrifuging at 6500 rpm for 5 min, the pellet was then resuspended in staining solution [50μM Tris pH 7.5, 1μM SYTOX green (Molecular Probes #S7020) and 1% RNase cocktail] and incubated 30 min at room temperature in the dark. Flow cytometry analysis was performed using a BD-Canto flow cytometer. The cytometer was run in linear mode, calibrated to the 1N population of the RH parasite reference control, and 70,000 events collected per sample. Histograms were generated using FACSDiva software.

#### Cell cycle IFA

Excysted Bradyzoites were cytospun (500 rpm for 5 min medium acceleration) onto slides, then co-stained for IMC1 and genomic DNA (DAPI) or separately co-stained for SRS9 and SAG1. IFA analysis of ME49EW Day 2-, 7- and 9-recrudescent populations were also analyzed by co-staining with antibodies against IMC1 and centrin and DAPI (genomic DNA). Parasite vacuoles were quantified for single (G1 phase) versus double (S/M phases) centrosomes, and also for the presence of internal daughters in 50 randomly selected vacuoles x3 for each sample.

#### Hoechst live stain

Purified cysts were stained following brain tissue homogenization using a 1:4 dilution of 12.3mg/ml of Hoechst stain (Thermo Scientific) and incubated for 20-30 minutes prior to being viewed on a Zeiss Axiovert microscope, at 400x magnification and images collected using Zeiss Zen software.

## ACKNOWLEDGEMENTS

*Toxoplasma gondii* genomic and RNA-sequencing data were accessed via http://ToxoDB.org. The authors thank Drs. William Sullivan Jr. and Krista Brooks for their insights and contributions that helped this paper come together. We thank Drs. John Boothroyd, Louis Weiss, and Gary Ward for antibodies used in this study. Tissue cyst RNA-sequencing gene data was generated by the Moffitt Cancer Center Molecular Genomics core facility. This study was supported by NIH NIAID R01 grants AI124682 to MWW and EHW, AI122760 to MWW and DA048815 to EHW.

